# Structure-Guided Optimization and Functional Characterization of Small Molecule Antagonists Targeting CD28 Costimulation

**DOI:** 10.1101/2025.10.06.680836

**Authors:** Saurabh Upadhyay, Hossam Nada, Sungwoo Cho, Moustafa T. Gabr

**Author notes:** **Corresponding author:** Dr. Moustafa T. Gabr.

## Abstract

CD28 is the prototypical costimulatory receptor that integrates with TCR signaling to sustain T-cell activation, proliferation, and survival. While indispensable for adaptive immunity, persistent CD28 signaling drives autoimmunity, graft-versus-host disease, and inflammatory pathology. Despite its therapeutic relevance, CD28 has long been regarded as an undruggable target due to its flat, solvent-exposed dimer interface, restricting modulation to biologics. Here, we describe a structure–activity relationship (SAR) campaign to optimize a small molecule CD28 inhibitor. Guided by biophysical profiling and functional assays, derivatives of the 8VS and 22VS chemotypes were evaluated, leading to the identification of BPU11 as a chemically tractable lead with improved pharmacokinetic stability, aqueous solubility, and plasma persistence. BPU11 consistently disrupted CD28–B7 interactions across biochemical and cellular systems, and potently suppressed T-cell activation in both a tumor–PBMC co-culture and a human PBMC– mucosal tissue model, functionally mimicking the biologic antagonist FR104. Molecular docking and dynamics simulations revealed engagement of the lipophilic canyon of CD28 through stabilizing hydrogen-bonding and hydrophobic interactions. These findings expand the pharmacological space of immune checkpoint blockade beyond antibodies and position BPU11 as a foundation for next-generation immunotherapies.

## 1. Introduction

Therapeutic antibodies directed against immune checkpoints have transformed clinical oncology, yet their impact is tempered by incomplete patient responses, relapse, and immune-related toxicities (Zamani & Sacha, 2025). While PD-1 and CTLA-4 remain the most clinically advanced checkpoints, accumulating evidence underscores the central role of costimulatory receptors in shaping T-cell immunity (Buchbinder & Desai, 2016). Among these, CD28 is the prototypical costimulatory receptor, constitutively expressed on naïve T cells and essential for full activation, IL-2 secretion, and clonal expansion (S. Upadhyay, B. Kaur, et al., 2025). Its ligation with CD80 (B7-1) and CD86 (B7-2) on antigen-presenting cells integrates with TCR signaling to reinforce effector functions, metabolic fitness, and survival (Levine et al., 1995). Beyond initial activation, CD28 also sustains T-cell responses in chronic antigen exposure, including within tumors, where it can maintain proliferation even under PD-1–mediated inhibition (Humblin et al., 2023). Conversely, unchecked CD28 signaling contributes to pathological immune activation, manifesting in autoimmunity, graft-versus-host disease, and chronic inflammation, thereby positioning the CD28–B7 axis as a therapeutic target of dual relevance in both oncology and immunopathology (Orvain et al., 2022; Porciello et al., 2018).

Attempts to modulate CD28 have largely relied on biologics (Crepeau & Ford, 2017). The catastrophic cytokine storm observed with the CD28 superagonist TGN1412 highlighted the risks of indiscriminate activation (Suntharalingam et al., 2006), while subsequent development of antagonistic antibodies (e.g., FR104, lulizumab) and CTLA-4-Ig fusion proteins (abatacept, belatacept) provided proof of concept for selective inhibition (Poirier et al., 2015; Schwarz et al., 2016; Shi et al., 2017). However, these agents remain limited by systemic immunosuppression, immunogenicity, and restricted tissue penetration inherent to macromolecular therapies (Chames et al., 2009). Small-molecule inhibitors could address these gaps, offering oral bioavailability, improved tumor penetration, and reversible pharmacodynamics (Zhang et al., 2025). Historically, CD28 has been considered a poor candidate for such an approach due to its flat, solvent-exposed binding interface (Calvo-Barreiro, Upadhyay, et al., 2025; S. Upadhyay, S. Cho, et al., 2025). Yet advances in molecular dynamics simulations, cryptic pocket mapping, and pharmacophore modeling have revealed druggable opportunities within the CD28 dimer interface, opening the possibility of rationally designed small-molecule modulators (Nada et al., 2025; Saurabh Upadhyay et al., 2025). These developments place CD28 at the forefront of next-generation checkpoint modulation strategies, not as a replacement but as a complement to existing PD-1/CTLA-4 blockade. Notably, this strategy aligns with a broader trend in drug discovery where challenging protein–protein interfaces and atypical binding sites—such as those in PKM2(S. Upadhyay, M. Bhardwaj, et al., 2025), G protein–coupled receptors (GPCRs)(Nishimura et al., 2020), and D-amino acid oxidase (DAAO) (Dave et al., 2023; Khan et al., 2024)—are being successfully exploited using structure-guided design and dynamic binding site mapping. As interest grows in fine-tuning immune responses through PPI blockade, our findings offer timely and translationally relevant insights into next generation immunomodulators.

In our recent structure-guided discovery effort (Saurabh Upadhyay et al., 2025), we reported the to our knowledge, first validated small-molecule antagonists of CD28. Among them, compound 22VS emerged as a functional lead, demonstrating reproducible binding and submicromolar inhibition of CD28–B7 interactions across biochemical and translational assays. In parallel, we also identified 8VS as a scaffold with consistent biophysical binding and functional inhibition. To assess the optimization potential of both chemotypes, we undertook a structure activity relationship (SAR)-by-catalogue approach to procure derivatives of 22VS and 8VS.

Our optimization campaign yielded three optimized derivatives—BPU11, BPU16, and BPU18— that preserve the parent pharmacophore while introducing modifications that enhance CD28 engagement. These compounds not only validate the tractability of the parent compounds for scaffold-based optimization but also expand the chemical space of CD28 inhibitors, providing a foundation for the development of next-generation small-molecule immunomodulators.

## 2. Materials and Methods

### 2.1 Screening of Small-Molecule Binders to Human CD28 Using Dianthus TRIC Assay

His-tagged human CD28 protein (Acro Biosystems) was labeled with RED-tris-NTA 2nd Generation dye (NanoTemper Technologies, Cat. #MO-L018) following the manufacturer’s protocol. A labeling mixture containing 20 nM dye and 40 nM CD28 was prepared in assay buffer [PBS, pH 7.0, containing 0.005% Tween-20] and incubated for 30 min at room temperature in the dark.

For high-throughput screening, a focused library of 40 catalogue-derived compounds (19 analogues each from the 8VS and 22VS chemotypes; Enamine screening collection) was tested. Labeled CD28 was mixed 1:1 with each compound (final concentration 100 µM in 2% DMSO) or control solutions and incubated for 15 min at room temperature. Human CD80 (2 µM) served as a positive control, and 2% DMSO was used as a negative control. After centrifugation (1000 × g, 1 min), 20 µL samples were loaded into the Dianthus NT.23 Pico instrument (NanoTemper Technologies) and analyzed at 25 °C. Fluorescence at 670 nm was recorded for 5s before and 5s after IR-laser activation. Normalized TRIC signals (Fnorm) were calculated as Fhot/Fcold. Each compound was tested in three technical replicates, with positive and negative controls included in duplicate at regular intervals across the 384-well plate.

### 2.2 Microscale Thermophoresis (MST) for Binding Affinity

For binding affinity validation, His-tagged CD28 protein (Acro Biosystems) was labeled with RED-tris-NTA 2nd Generation dye (NanoTemper Technologies) using the Monolith His-Tag Labeling Kit according to the manufacturer’s protocol. A solution of 50 nM dye and 100 nM CD28 was prepared in assay buffer [PBS, pH 7.0, containing 0.005% Tween-20] and incubated for 30 min at room temperature in the dark.

Compounds were prepared as 12-point, 1:2 serial dilutions starting from 500 µM in assay buffer containing 2% DMSO. Labeled protein and compound solutions were mixed 1:1 (final volume 15 µL) and incubated for 15 min at room temperature in the dark. Samples were loaded into Monolith NT.115 Premium Capillaries (NanoTemper Technologies, Cat. #MO-K022) and measured at 25 °C using the Monolith NT.115 instrument with 40% excitation power and medium MST power.

Normalized fluorescence (Fnorm) was calculated as the ratio of post-laser to pre-laser fluorescence. Each compound was tested in four technical replicates, and confirmed binders were validated in three independent biological experiments. Dissociation constants (Kd) were determined by fitting dose–response curves using the MO.Affinity Analysis software and GraphPad Prism 10 (four-parameter nonlinear regression).

### 2.3 Evaluation of CD28–CD80 Binding Inhibition via Competitive ELISA

Competitive ELISA assays were performed using the CD28:B7-1 [Biotinylated] Inhibitor Screening Kit (BPS Bioscience, Cat. #72007), following the manufacturer’s instructions with minor modifications. Briefly, 96-well plates were coated with recombinant human CD28 (2 µg/mL in PBS, 50 µL/well) overnight at 4 °C. The next day, wells were washed with 1× Immuno Buffer and blocked with Blocking Buffer for 1 h at room temperature.

Serial dilutions of test compounds were co-incubated with biotinylated CD80 (5 ng/µL) for 1 h at room temperature to allow competitive binding to immobilized CD28. Wells without coating served as ligand controls, while wells incubated with inhibitor buffer instead of compound served as negative controls. After washing, plates were incubated with streptavidin–HRP (1:1000 in Blocking Buffer) for 1 h. Chemiluminescent substrate was added, and luminescence was immediately measured using an Infinite M1000 Pro Microplate Reader (Tecan, Switzerland). IC_50_ values were determined by fitting concentration–response curves using a four-parameter logistic regression in GraphPad Prism 10. All experiments were performed in triplicate.

### 2.4. Molecular Docking

Molecular docking studies were performed using Schrödinger Suite 2023.4 to investigate the binding mode of BPU11 with the CD28 protein. The crystal structure of CD28 was obtained from the PDB (PDB ID: 1YJD1)(Evans et al., 2005) and prepared using the Protein Preparation Wizard, which included optimization of hydrogen bond networks, removal of crystallographic water molecules, and energy minimization using the OPLS4 force field. The ligand BPU11 was prepared using LigPrep wizard at default settings. Grid generation was centered on the conserved binding region comprising residues 99-104, which has been previously identified as the primary ligand binding site shared between CD28 and CTLA-4 (Calvo-Barreiro, Nada, et al., 2025). Docking calculations were performed using Glide in standard precision (SP) mode, with a maximum of 10 poses per ligand retained for analysis.

### 2.5 Molecular Dynamics Simulations

Molecular dynamics (MD) simulations were conducted using GROMACS 2024.4 to evaluate the stability and dynamic behavior of the BPU11/CD28 complex. The CHARMM27 all-atom force field was employed for the MD simulations. The complex was solvated in a cubic box with TIP3P water molecules, maintaining a minimum distance of 1.0 nm between the protein and box edges. The system was neutralized with appropriate counter-ions and subjected to energy minimization using the steepest descent algorithm. Following minimization, the system underwent equilibration in two phases: NVT equilibration at 300 K for 100 ps using the V-rescale thermostat, followed by NPT equilibration at 1 bar for 100 ps using the Parrinello-Rahman barostat. Production MD simulations were performed for 50 ns with a 2 fs time step, saving coordinates every 10 ps for subsequent analysis.

### 2.6 Cell Viability Assay

Cytotoxicity of test compounds was assessed using the MTS assay (CellTiter 96® AQueous One Solution, Promega). Jurkat T cells were seeded in 96-well plates and cultured in complete RPMI medium with 10% FBS for 24 h to reach ∼40% confluence. Cells were then treated with serial dilutions of compounds (10–500 µM) or vehicle controls (0.1% DMSO, 0.1% Tween-20) for 24 h. MTS reagent was added according to the manufacturer’s protocol, and absorbance was measured at 490 nm using an Infinite M1000 Pro Microplate Reader. Cell viability was expressed as a percentage of vehicle-treated controls. Each experiment was performed in triplicate, with results representing the mean ± SEM from at least three independent biological replicates.

### 2.7 NanoBit Luciferase Complementation Assay

Protein–protein interactions between CD28 and its ligands were evaluated using the NanoBit luciferase complementation assay (Promega). CHO-K1 cells were transfected with plasmids encoding CD28 fused to SmBiT and either CD80 or CD86 fused to LgBiT. Transfected cells were seeded into white 96-well plates and cultured until ∼70% confluence.

Cells were washed twice with PBS, then incubated in Hank’s Balanced Salt Solution (HBSS) containing test compounds (0.1–500 µM) for 2 h at 37 °C. After treatment, furimazine substrate (10 µM final concentration) was added and incubated for 10 min at room temperature. Luminescence was measured using an Infinite M1000 Pro Microplate Reader. Relative luminescence units (RLU) were normalized to DMSO-treated controls after background subtraction.

All experiments were performed in triplicate, and data represent mean ± SEM from at least three independent experiments.

### 2.8 CD28 Blockade Bioassay

The CD28 Blockade Bioassay (Promega, Cat. #JA6101) was used to assess functional inhibition of CD28 signaling. Jurkat CD28 Effector Cells (2 × 10⁴ cells/well) were seeded in white 96-well plates and pre-incubated for 5 min with serial dilutions of test compounds (10-point, 1:1 dilution series starting at 200 µM, final 1% DMSO). Anti-CD28 control antibody (Promega, Cat. #K1231) was included as a positive control. After pre-incubation, aAPC/Raji Cells (2 × 10⁴ cells/well) were added, and plates were incubated for 5 h at 37 °C, 5% CO_2_. Bio-Glo™ Luciferase Reagent (Promega) was then added, and luminescence was measured on a GloMax® Discover System (Promega). Dose–response curves were generated using GraphPad Prism 10 with a four-parameter logistic regression model to calculate IC_50_ values. All experiments were performed in triplicate, and results represent mean ± SEM. The assay demonstrated tolerance to up to 10% pooled human serum, confirming robustness under physiological conditions.

### 2.9 Pharmacokinetic profiling

The preliminary evaluation of PK parameters for BPU11 was performed as previously reported by us (Abdel-Rahman et al., 2025).

### 2.10 Evaluation of T Cell Activation in Tumor-PBMC Coculture

To assess the dose-dependent impact of CD28 inhibition on T cell activation in a translational human system, we employed a 3D tumor–PBMC co-culture assay. Tumor spheroids were generated by seeding A549 (or MDA-MB-231) cells in ultra-low attachment 96-well plates. After 48 hours, spheroids were treated with recombinant human interferon-gamma (IFN-γ, 50 ng/mL) for 24 hours. Freshly thawed human peripheral blood mononuclear cells (PBMCs) were added at a 5:1 effector-to-target (E:T) ratio, and co-cultures were stimulated with soluble anti-CD3 antibody (0.3 μg/mL, clone OKT3) to model TCR activation.

The tested compound (BPU11) was added at the initiation of co-culture. FR104 (10 μg/mL), a reference anti-CD28 Fab’ biologic, served as a positive control. BPU11 was evaluated at three concentrations (5, 10, and 25 μM). Vehicle and anti-CD3–only conditions were included as controls. After 48 hours, supernatants were harvested and analyzed for IFN-γ and IL-2 levels using human-specific ELISA kits (BioLegend). Soluble CD69 (sCD69), released from activated T cells, was quantified using a human CD69 ELISA kit (ThermoFisher).

### 2.11 CD28-Dependent Cytokine Release in a Human PBMC–Mucosal Co-culture Model

To evaluate the immunomodulatory activity of BPU11 in a mucosal-relevant human immune interface, we employed a commercially available human airway epithelial tissue model (MucilAir™, Epithelix) co-cultured with PBMCs. Cryopreserved human PBMCs (StemCell Technologies) were thawed and rested for 4 hours in RPMI-1640 medium supplemented with 10% heat-inactivated human AB serum, 2 mM L-glutamine, and 1% penicillin–streptomycin.

MucilAir™ inserts were equilibrated in the manufacturer’s proprietary maintenance medium for 24 hours at 37 °C and 5% CO2. PBMCs were resuspended at 2 × 10^6^ cells/mL and overlaid (200 µL per insert) onto the apical surface of the MucilAir™ tissue in 24-well Transwell plates. Co-cultures were stimulated with plate-bound anti-CD3 (1 µg/mL) and soluble anti-CD28 (1 µg/mL) in the presence of: vehicle control (0.1% DMSO), BPU11 (5, 10, or 25 µM), or FR104 (10 µg/mL; anti-CD28 Fab’ fragment).

After 48 hours, apical supernatants were harvested and analyzed for IFN-γ, IL-2, and TNF-α concentrations using human ELISA kits (BioLegend) according to manufacturer instructions. Cytokine concentrations were interpolated from standard curves and expressed in pg/mL. All conditions were performed in triplicate (three independent experiments).

## 3. Results

### 3.1 Biophysical Screening and Affinity Profiling Using TRIC and MST

To assess the optimization potential of the parent scaffolds (8VS and 22VS), we implemented a two-stage biophysical pipeline integrating high-throughput TRIC-based (Dianthus) screening with orthogonal validation by microscale thermophoresis (MST). A focused library of 40 catalogue-derived analogues (Table S1, Supporting Information), comprising derivatives each from the 8VS and 22VS chemotypes was procured from Enamine.

All derivatives (BPU1–BPU40, Table S1) incorporate a conserved 1,3,4-thiadiazole–triazole– thioamide pharmacophore, which serves as the core recognition motif for CD28 binding. Structural diversification was introduced along three key moieties to explore the relationship between chemical modifications and biological activity. First, a broad range of aryl and heteroaryl terminal substitutions was introduced at the triazole-thioamide terminus to modulate hydrophobic and π–π interactions within the lipophilic canyon of CD28. These include electron-donating (–Me, –OMe) and electron-withdrawing (–Cl, –F, –CF_3_, –NO_2_) substituents, as well as heteroaromatic replacements that altered electronic density and hydrogen-bonding potential. Second, the central heteroaryl scaffold was varied to adjust conformational rigidity and polarity, influencing hydrogen-bonding capacity and alignment within the CD28 binding groove. These subtle core changes allowed evaluation of how planarity and ring heteroatom placement affect receptor complementarity. Third, terminal linkers and aliphatic fragments were modified to yield compounds with variable lipophilicity and steric bulk. Collectively, these modifications generated a chemically diverse library aiming to guide future optimization of small molecule CD28 antagonists.

The 40 compounds were screened at 100 μM under standard buffer conditions (PBS, pH 7.0, 0.005% Tween-20, 4% DMSO). Each compound was tested in triplicate, and 40 negative controls were included for plate normalization. Analysis of normalized fluorescence (Fnorm) identified eight hits—BPU1, BPU7, BPU11, BPU16, BPU18, BPU28, BPU37, and BPU38—that exceeded the ±5 SD threshold relative to controls (Figure 1A). These candidates were subsequently evaluated by concentration-dependent MST. Three analogues, BPU11, BPU16, and BPU18 (Table 1), displayed reproducible sigmoidal binding curves with dissociation constants (Kd) of 6.5 ± 2.8 µM, 57.8 ± 9.8 µM, and 30.5 ± 4.8 µM, respectively, confirming specific CD28 engagement (Figure 1B–D). (Figure 1B–D). Remarkably, the optimized analogue BPU11 exhibited the strongest CD28 affinity (Kd = 6.5 ± 2.8 µM), representing a nearly five-fold improvement over the parent scaffold 8VS (Kd = 29.57 ± 12.69 µM, Table 1) and an eight-fold enhancement relative to 22VS (Kd = 52.45 ± 18.69 µM, Table 1).

**Figure 1.**
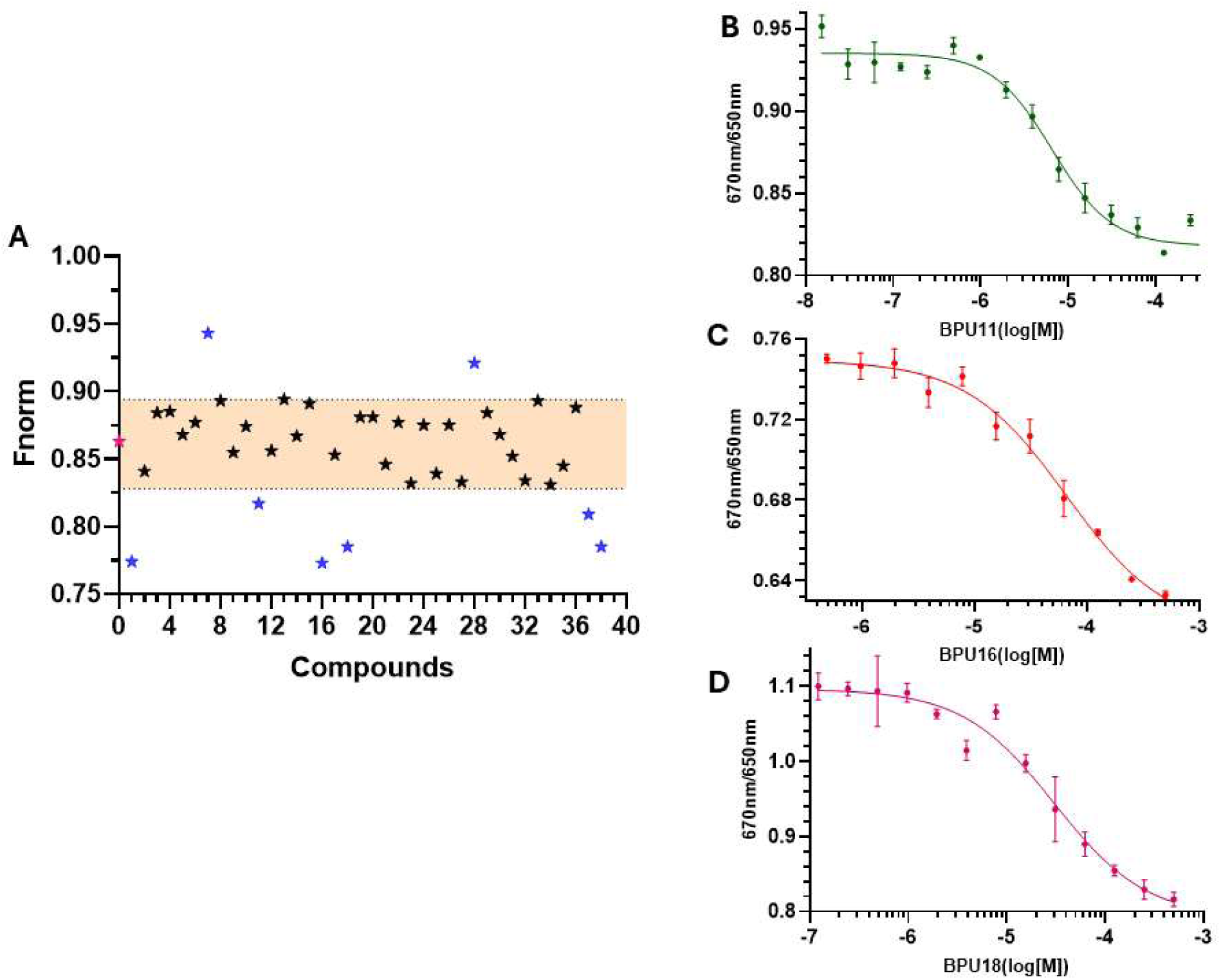
Biophysical identification and validation of SAR-derived CD28 binders. (A) TRIC-based Dianthus screen of 40 catalogue-derived analogues (20 each from 22VS and 8VS chemotypes). Normalized fluorescence (Fnorm) values at 100 μM identified eight compounds exceeding the ±5 SD threshold (shaded region). (B–D) MST validation of three SAR-derived hits. Sigmoidal dose–response binding curves confirmed specific CD28 engagement with dissociation constants of 6.5 ± 2.8 μM for BPU11 (B), 57.8 ± 9.8 μM for BPU16 (C), and 30.5 ± 4.8 μM for BPU18 (D). Data represent mean ± SEM from three independent experiments, fitted using a four-parameter nonlinear regression model.

**Table 1.**
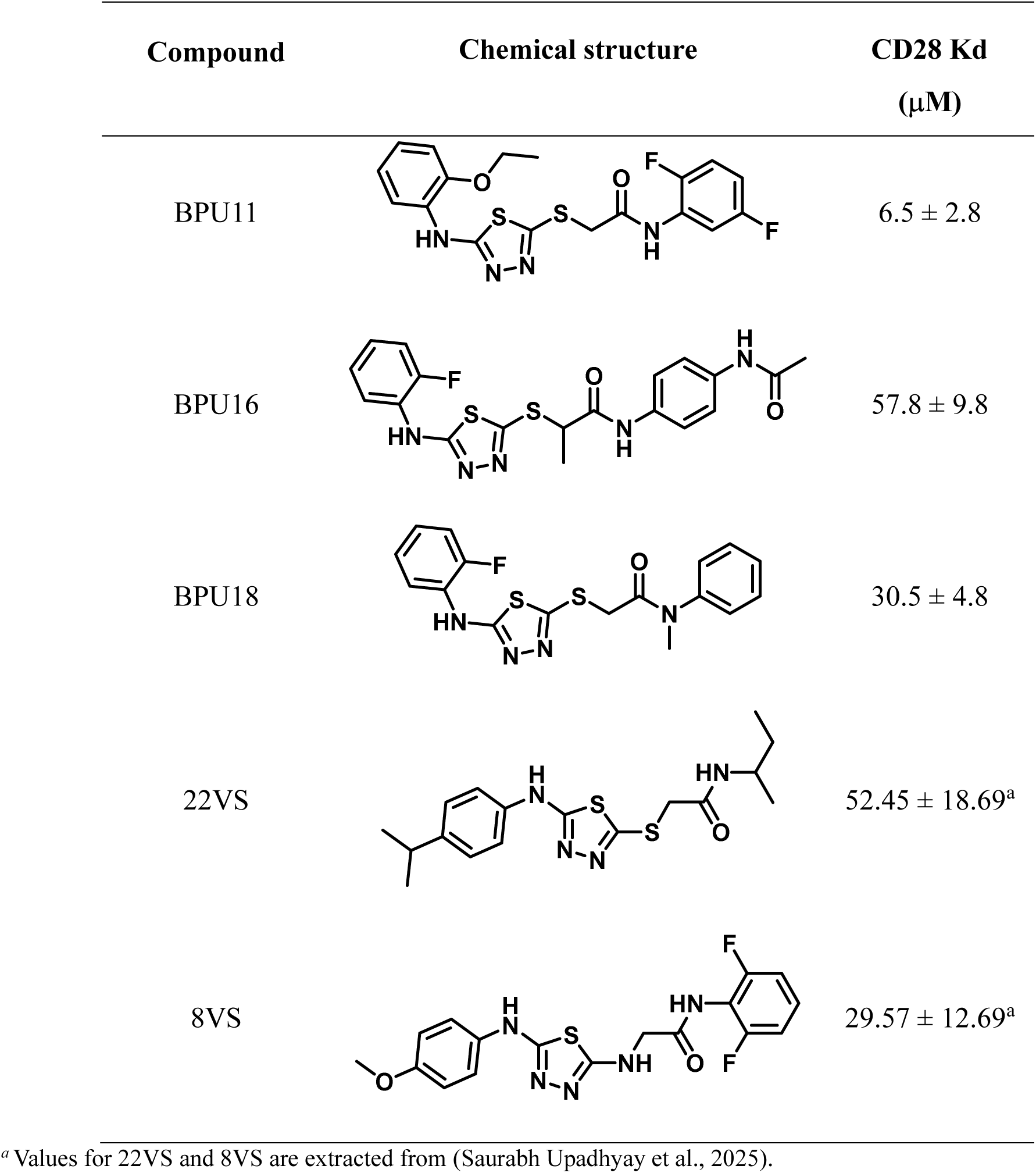
Chemical structures of the top optimized leads and parent hit compounds, together with their CD28 binding affinities determined by MST.

| Compound | Chemical structure | CD28 Kd<br>($\mu\text{M}$) |
| --- | --- | --- |
| BPU11 | | $6.5 \pm 2.8$ |
| BPU16 | | $57.8 \pm 9.8$ |
| BPU18 | | $30.5 \pm 4.8$ |
| 22VS | | $52.45 \pm 18.69^a$ |
| 8VS | | $29.57 \pm 12.69^a$ |

<sup>a</sup> Values for 22VS and 8VS are extracted from (Saurabh Upadhyay et al., 2025).

### 3.2 Functional Inhibition of CD28–CD80 Interaction in ELISA

The ability of the derivatives to disrupt CD28–ligand binding was evaluated using an ELISA-based CD28:B7-1 assay. Recombinant CD28 was immobilized on 96-well plates and incubated with biotinylated CD80 in the presence of serial dilutions of test compounds. Bound ligand was quantified by streptavidin–HRP detection, and inhibition curves were fitted from three independent experiments.

All compounds inhibited CD28–CD80 binding in a dose-dependent manner, producing sigmoidal concentration–response curves (Figure 2A–D). The parent scaffold 8VS exhibited modest activity with an IC_50_ of 71.61 μM. Among the derivatives, BPU11 displayed the strongest inhibition, with an IC_50_ of 18.89 μM, representing a nearly four-fold improvement over the parent scaffold. BPU18 also showed enhanced potency with an IC_50_ of 45.16 μM, whereas BPU16 was weaker (IC_50_ = 86.74 μM).

**Figure 2.**
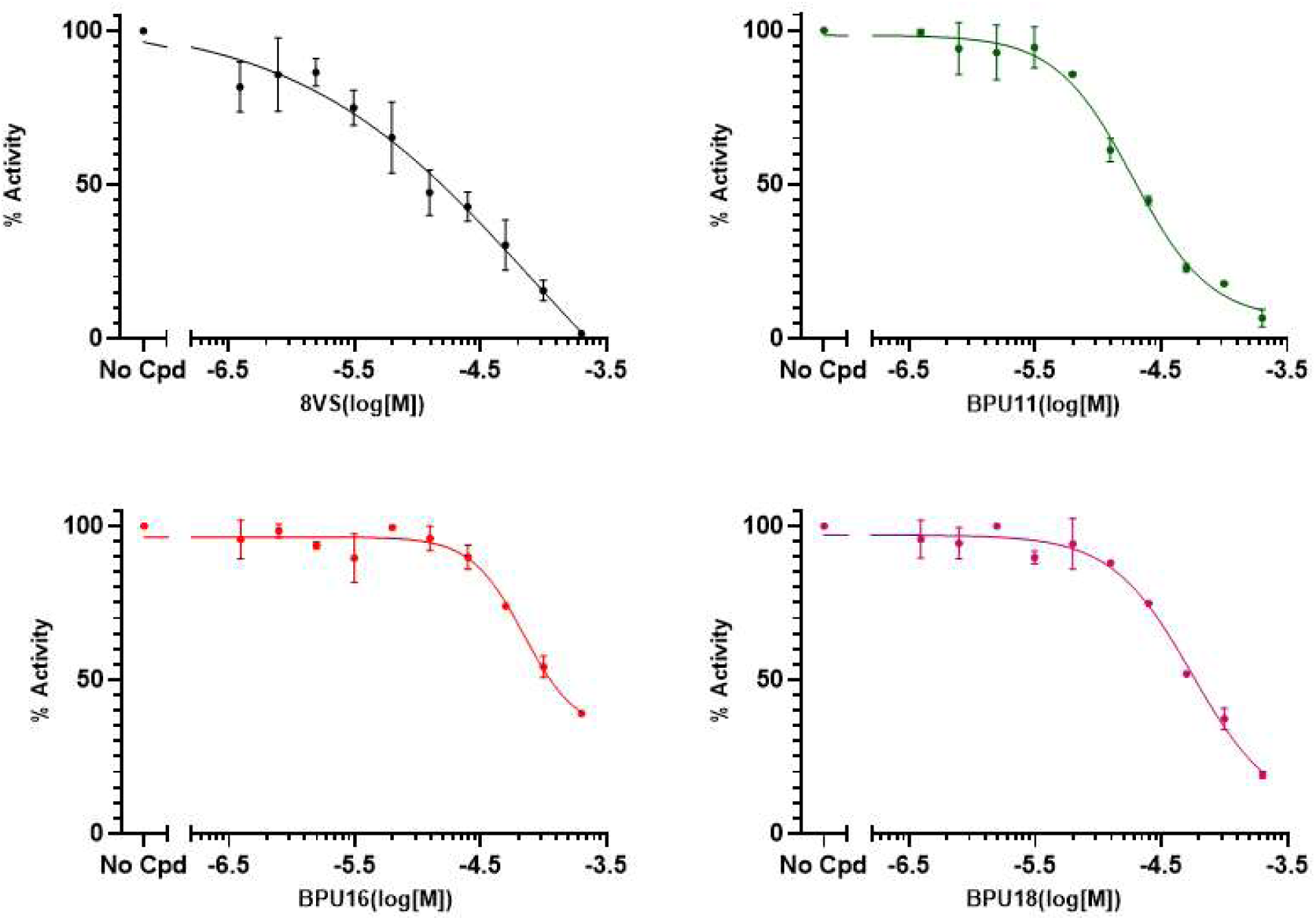
Functional inhibition of CD28–CD80 interaction by 8VS and its derivatives. ELISA-based CD28:B7-1 binding assays were performed using recombinant CD28 immobilized on 96-well plates and biotinylated CD80 as ligand. Increasing concentrations of test compounds were added, and binding was detected with streptavidin–HRP. All compounds inhibited CD28– CD80 binding in a concentration-dependent manner. Dose–response curves yielded IC_50_ values of 71.61 μM for 8VS (A), 18.89 μM for BPU11 (B), 86.74 μM for BPU16 (C), and 45.16 μM for BPU18 (D). Data represent mean ± SEM from at least three independent experiments, with nonlinear regression fits generated using a four-parameter logistic model.

The relative rank order of activity (BPU11 > BPU18 > 8VS > BPU16) closely paralleled the binding affinities determined by MST, reinforcing that the inhibitory effects reflect specific disruption of the CD28–B7 interaction.

### 3.3 Binding Mechanism of BPU11 to CD28 Revealed by Docking and Molecular Dynamics

The molecular docking results revealed that BPU11 bound to the CD28 protein at the conserved ligand binding site (Figure 3A). The docked pose shows the compound adopting a favorable conformation within the binding pocket, with the ligand making multiple interactions with key residues in the binding region. The 2D interaction diagram (Figure 3B) illustrates the detailed binding mode, showing that BPU11 forms hydrogen bonds with critical residues including HIS38, PHE93, LYS95, LYS109, SER110, and ASN111. BPU11 was predicted to establish both hydrophobic and polar interactions which contributed to the overall binding affinity. The presence of aromatic rings in BPU11 allows for π-π stacking interactions with aromatic residues in the binding site, while polar functional groups form hydrogen bonds with surrounding amino acids.

**Figure 3.**
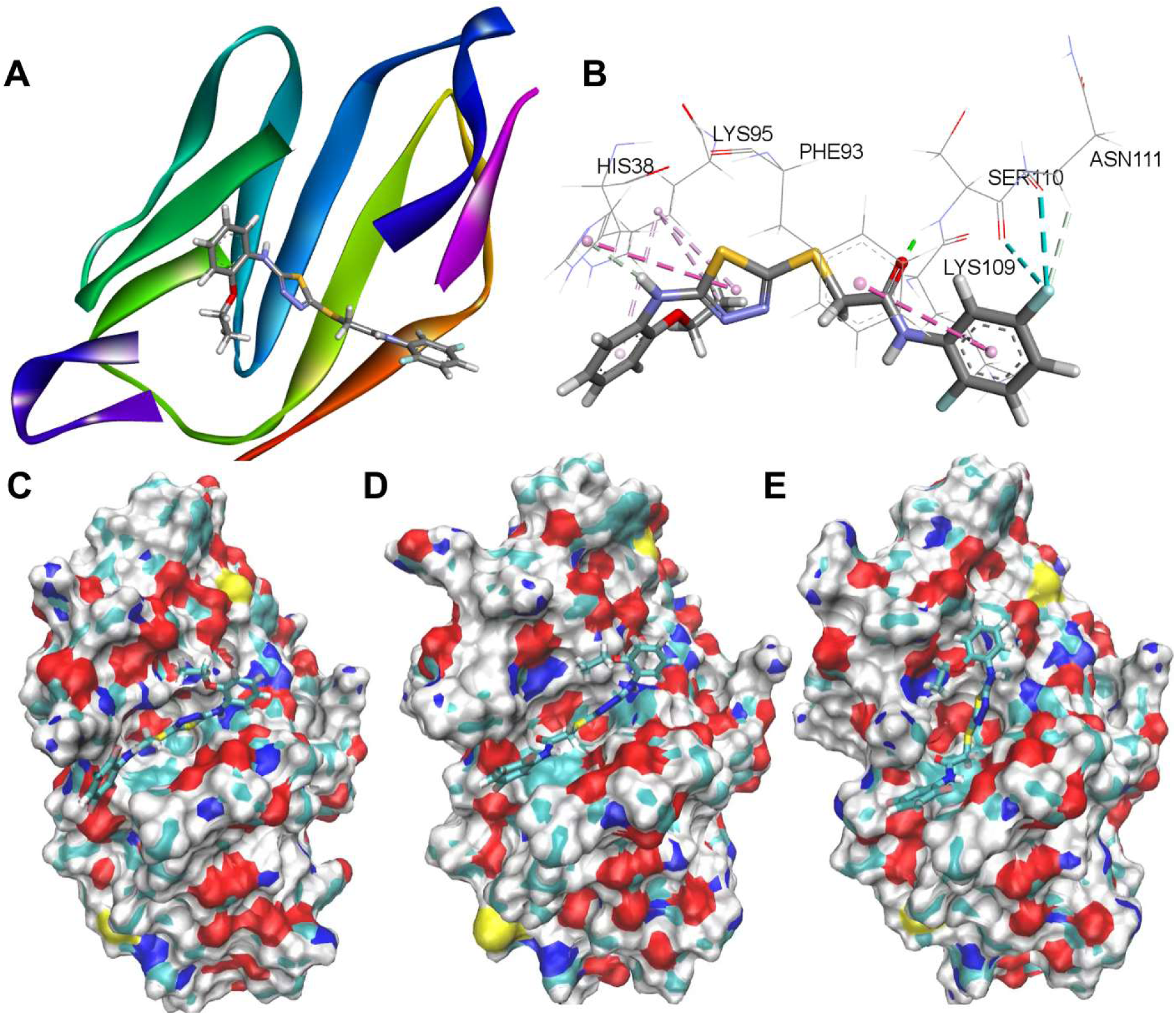
Structural analysis of BPU11 binding to CD28. (A) Three-dimensional representation of BPU11 (stick representation) docked within the CD28 binding site (ribbon representation, colored by secondary structure). The binding site is located within the conserved region comprising residues 99-104. (B) Two-dimensional interaction diagram showing detailed molecular interactions between BPU11 and CD28 residues, including hydrogen bonds (dashed lines) with key residues LYS95, PHE93, HIS38, SER110, LYS109, and ASN111. (C-E) Surface representations of the CD28 binding site showing conformational changes during molecular dynamics simulation.

Next, MD simulations were carried out to validate the results of the docking assay and assess the stability of the predicted complex. The MD simulation predicted dynamic stability for the BPU11/CD28 complex over the 50 ns simulation period. Root Mean Square Deviation (RMSD) analysis (Figure 4A) shows that the CD28-complex system (blue line) exhibits higher initial fluctuations compared to the unbound CD28 protein (green line) but reaches equilibrium after approximately 10 ns with an average RMSD of ∼0.5 nm. The unbound protein maintains lower RMSD values throughout the simulation (∼0.2 nm), indicating that ligand binding induces conformational changes that stabilize into a new equilibrium state.

**Figure 4.**
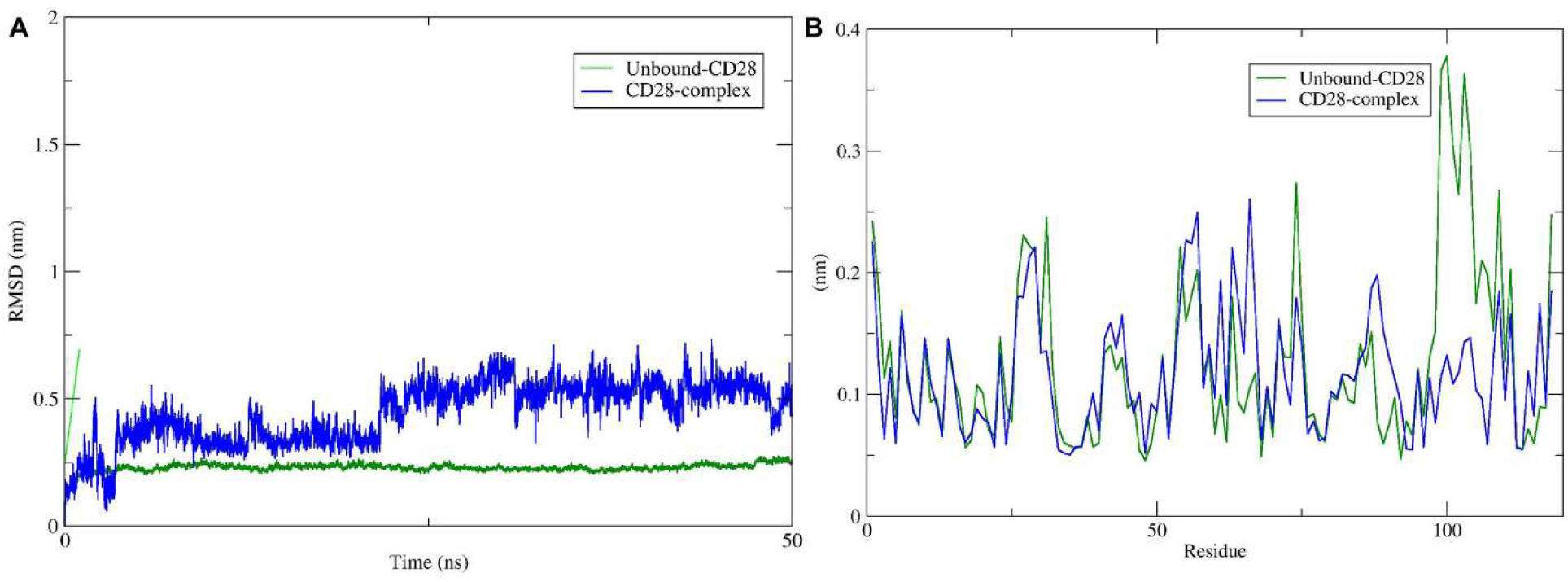
Root Mean Square Deviation (RMSD) and Root Mean Square Fluctuation (RMSF) analysis of the BPU11/CD28 complex. (A) RMSD trajectories over 50 ns simulation time comparing the BPU11/CD28 complex (blue) with unbound CD28 protein (green). The complex shows initial equilibration followed by stable dynamics with RMSD values around 0.5 nm. (B) Per-residue RMSF analysis showing differential flexibility patterns between bound (blue) and unbound (green) states, with notable changes in the 80-100 residue region indicating allosteric effects of ligand binding.

Root Mean Square Fluctuation (RMSF) analysis (Figure 4B) provides insights into the per-residue flexibility differences between bound and unbound states. The analysis reveals that certain regions (particularly around residues 80-100) show altered flexibility upon BPU11 binding, with some areas becoming more rigid while others show increased mobility. This suggests that ligand binding induces allosteric effects that propagate throughout the protein structure, potentially affecting its functional properties.

The conformational changes observed during the MD simulation (Figure 1C-E) illustrate the adaptive binding process. Initially, the binding site undergoes reorganization to accommodate the ligand, followed by gradual stabilization as favorable interactions are optimized. These induced-fit mechanisms are crucial for understanding how BPU11 achieves its binding specificity and affinity.

The Free Energy Landscape (FEL) analysis (Figure 5) constructed from the first two principal components of the MD trajectory reveals the conformational space explored by the BPU11/CD28 complex. The landscape shows three distinct stable conformational states, with one predominant basin indicating the most favored binding conformation. The depth of the primary energy minimum (shown in red, ∼16 kJ/mol) suggests strong stabilization of this conformation. The presence of alternative conformational states indicates some degree of flexibility in the binding mode, which may be important for the compound’s biological activity.

**Figure 5.**
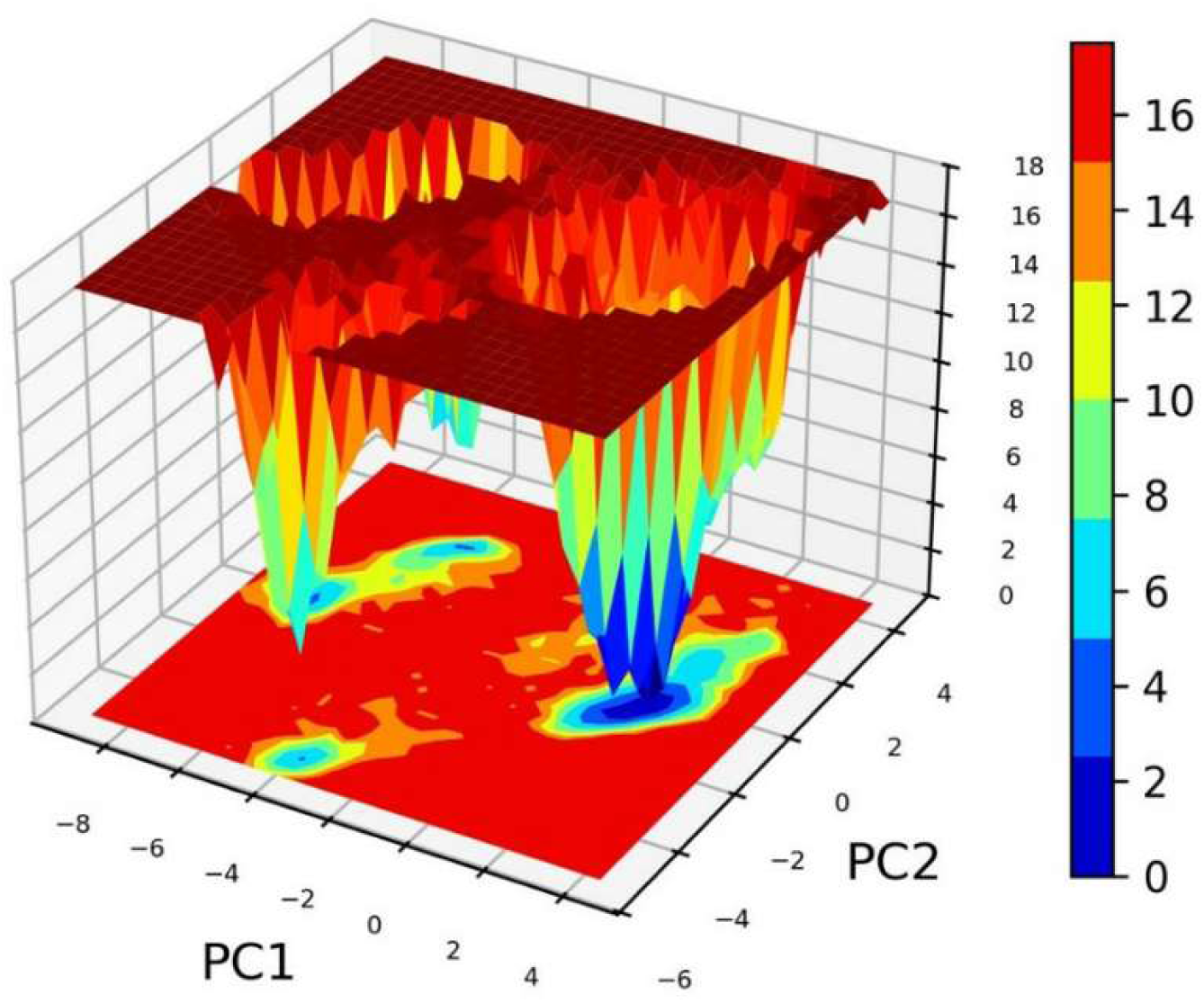
Free Energy Landscape (FEL) of the BPU11/CD28 complex. Three-dimensional representation of the conformational free energy surface constructed from the first two principal components (PC1 and PC2) of the MD trajectory. The color scale represents free energy in kJ/mol, with blue indicating higher energy states and red indicating lower energy, more stable conformations.

The energy barriers between different states are relatively low (2-4 kJ/mol), suggesting that transitions between conformations can occur readily at physiological temperatures. This conformational flexibility may contribute to the compound’s ability to maintain binding even in the presence of competitive ligands or allosteric modulators.

The combined docking and MD results provide valuable insights for the optimization of BPU11 as a CD28-targeting therapeutic agent. The identification of key binding residues suggests specific structural modifications that could enhance binding affinity. The dynamic analysis reveals that maintaining the observed hydrogen bonding patterns while optimizing hydrophobic interactions could lead to improved potency. Furthermore, the conformational flexibility observed in the FEL analysis indicates that rigid analogs might not be optimal, and that maintaining some degree of flexibility could be beneficial for binding kinetics.

### 3.4 Cellular Inhibition of CD28–B7 Interactions and Cytotoxicity Evaluation

The functional effects of 8VS derivatives on CD28–B7 interactions were examined using the NanoBit luciferase complementation assay in CHO-K1 cells co-expressing CD28-SmBiT with either CD80-LgBiT or CD86-LgBiT (Bromley et al., 2001; van der Merwe et al., 1997). Based on biophysical and biochemical profiling, BPU11, BPU16, and BPU18 were selected for evaluation. Preliminary cytotoxicity testing in Jurkat T cells revealed that BPU16 and BPU18 reduced cell viability at concentrations ≥30 μM, and these compounds were therefore excluded from NanoBit assays.

CHO-K1 cells were treated with increasing concentrations of BPU11 (0.1–500 μM), and luminescence was measured after 2 h. BPU11 inhibited both CD28–CD80 and CD28–CD86 interactions in a concentration-dependent manner (Figure 6A–B). Maximal inhibition reached ∼40% for CD28–CD80 and ∼40–45% for CD28–CD86. Nonlinear regression analysis yielded IC_50_ values of 21.88 μM for CD28–CD80 and 38.69 μM for CD28–CD86.

**Figure 6.**
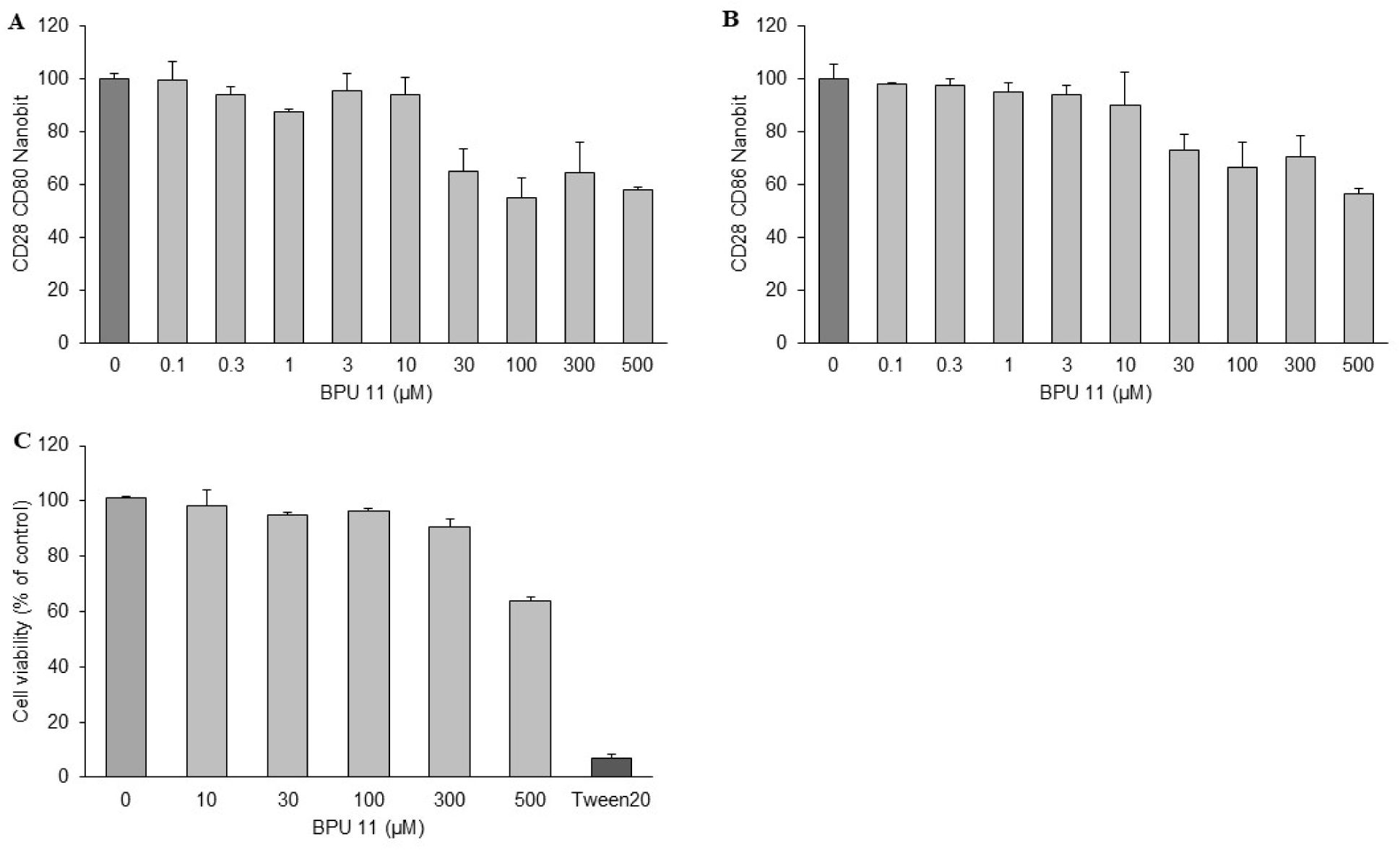
Cellular inhibition and cytotoxicity assessment of CD28–B7 interactions by BPU11. (A) Dose-dependent inhibition of CD28–CD80 interactions by BPU11 measured in CHO-K1 cells co-expressing CD28-SmBiT and CD80-LgBiT. (B) Dose-dependent inhibition of CD28–CD86 interactions by BPU11 in CHO-K1 cells co-expressing CD28-SmBiT and CD86-LgBiT. Cells were treated with BPU11 (0.1–500 μM) for 2 h at 37 °C, and luminescence was normalized to DMSO-treated controls. (C) Jurkat T-cell viability following exposure to BPU11 (10–500 μM) for 24 h, assessed by MTS assay. Viability remained >85% at concentrations up to 300 μM, with modest reductions (∼60% of control) at 500 μM. BPU16 and BPU18 were excluded from NanoBit functional assays due to cytotoxicity observed at ≥30 μM in preliminary viability screens. Data represent mean ± SEM from three independent experiments.

To confirm that inhibition was not attributable to nonspecific cytotoxicity, Jurkat T cells were exposed to BPU11 (10–500 μM) and viability was measured by MTS assay (Figure 6C). Cell viability remained >85% at concentrations up to 300 μM, with moderate reductions (∼60% of control) only at the highest concentration tested (500 μM).

### 3.5 Reporter-Based Functional Blockade Assay

The functional activity of 8VS derivatives was further examined using the CD28 Blockade Bioassay, a luciferase reporter system employing Jurkat effector cells co-cultured with Raji antigen-presenting cells. This assay measures inhibition of CD28-mediated costimulatory signaling in a cellular context.

Dose–response analysis demonstrated that both 8VS and BPU11 suppressed reporter activity in a concentration-dependent manner (Figure 7A–B). The parent scaffold 8VS exhibited moderate inhibition with an IC_50_ of 22.77 ± 6.8 μM. In comparison, BPU11 displayed enhanced activity, achieving an IC_50_ of 7.9 ± 2.3 μM, consistent with its stronger binding affinity and functional potency observed in earlier assays.

**Figure 7.**
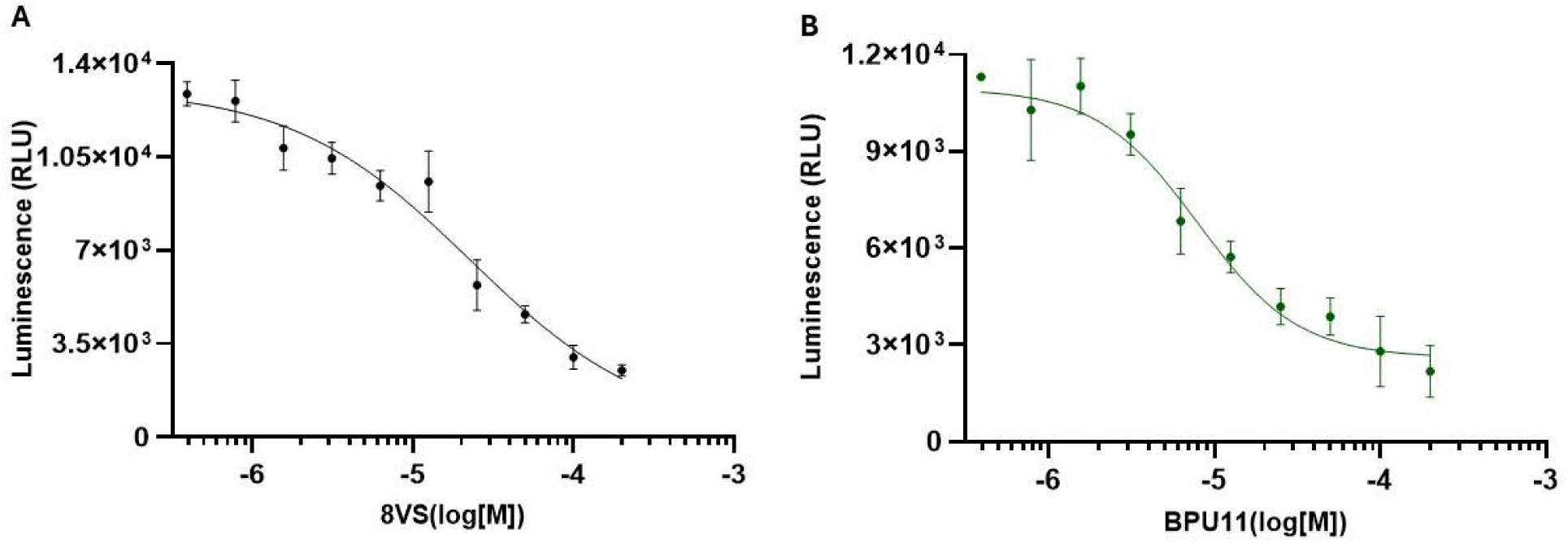
Reporter-based functional inhibition of CD28 signaling by 8VS and BPU11. Jurkat effector cells were co-cultured with Raji antigen-presenting cells in the CD28 Blockade Bioassay to measure inhibition of CD28-mediated costimulation. Cells were treated with increasing concentrations of compounds for 5 h, and luminescence was quantified using the Bio-Glo™ Luciferase Assay System. (A) 8VS inhibited CD28 signaling with an IC_50_ of 22.77 ± 6.8 μM. (B) BPU11 displayed stronger inhibition with an IC_50_ of 7.9 ± 2.3 μM. Data represent mean ± SEM from at least three independent experiments, and curves were fitted using a four-parameter logistic regression model.

### 3.6 Pharmacokinetic (PK) Profiling

To characterize the developability of the optimized CD28 inhibitor BPU11, we performed an in vitro pharmacokinetic (PK) assessment under conditions identical to those previously used for 22VS. The results are summarized in Table 2. BPU11 exhibited markedly enhanced solubility, achieving concentrations of 125 µM in PBS and 138 µM in FaSSIF, outperforming 22VS (93 µM and 108 µM, respectively). Despite a moderate increase in lipophilicity (LogD_7.4_ = 3.10), BPU11 maintained balanced physicochemical properties conducive to oral absorption. Its passive permeability measured by PAMPA (6.3 × 10⁻⁶ cm/s) confirmed efficient membrane diffusion.

**Table 2.**
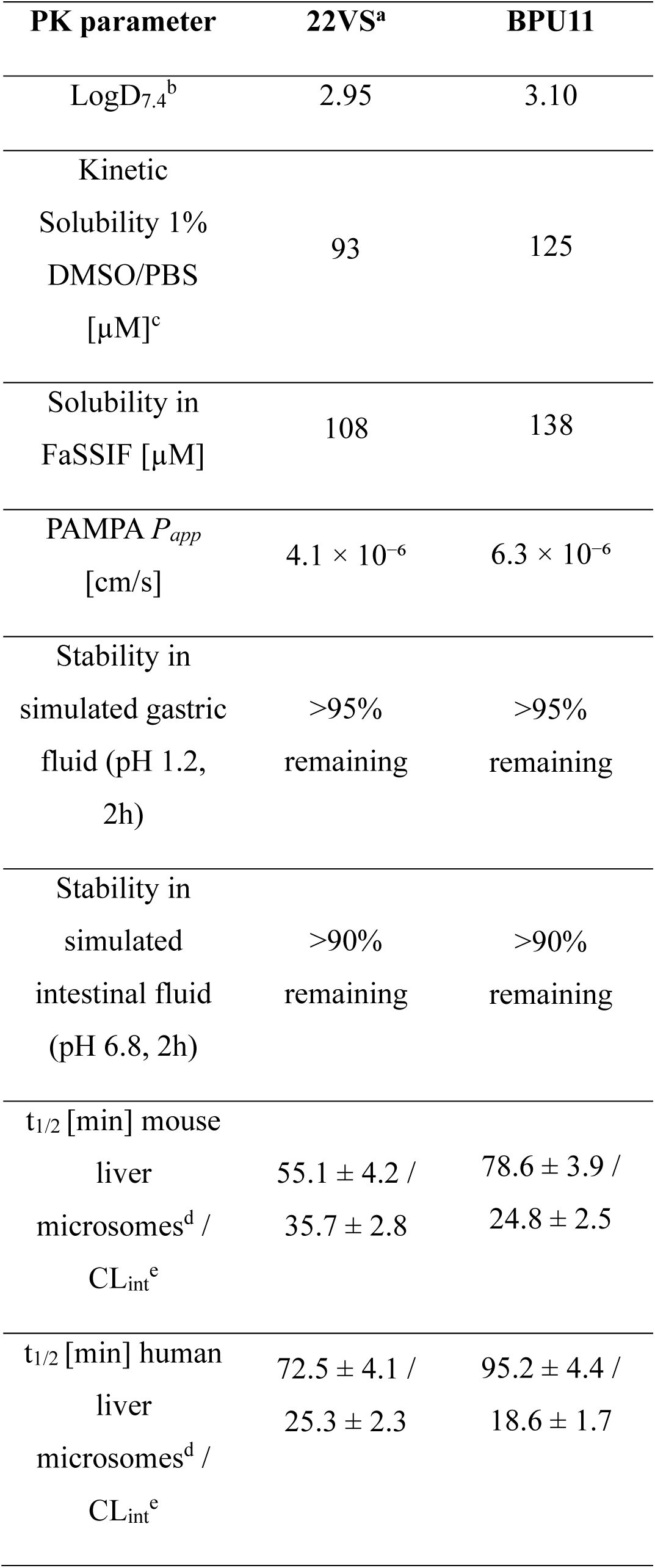
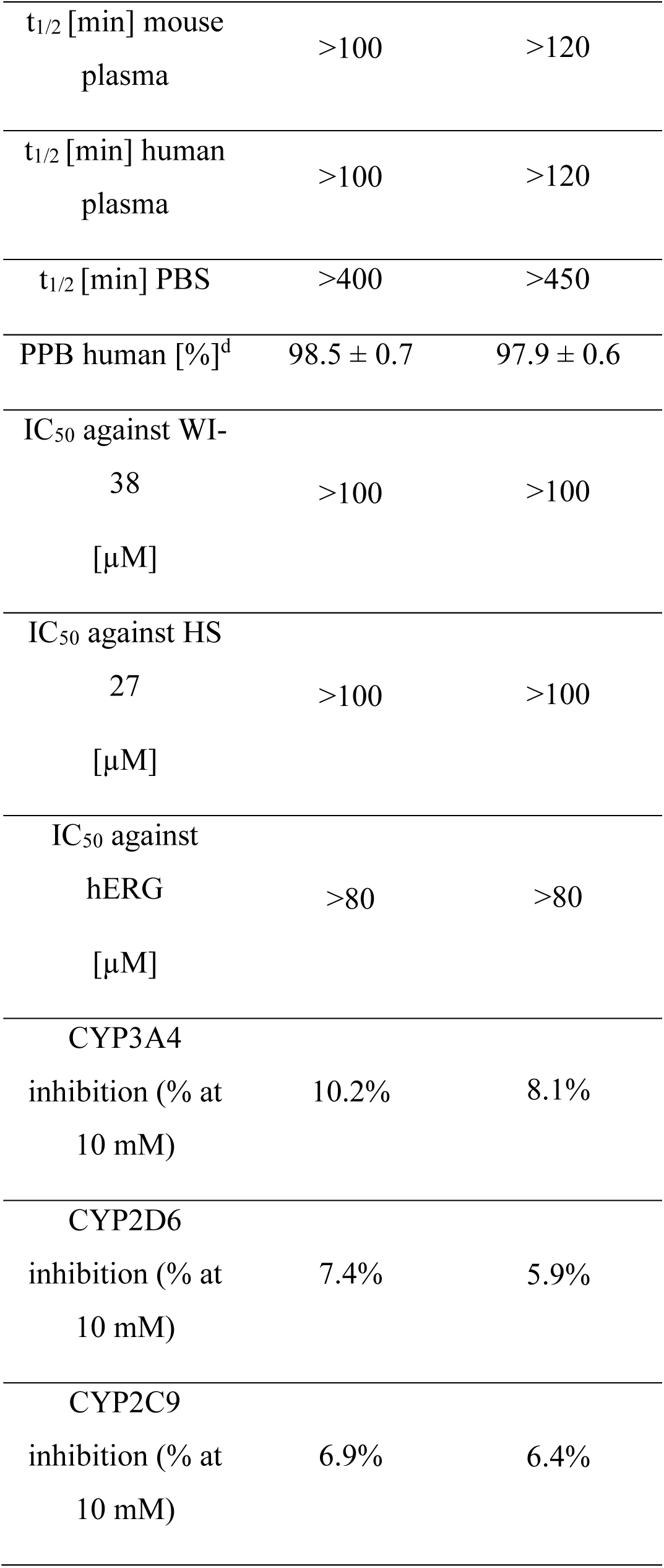
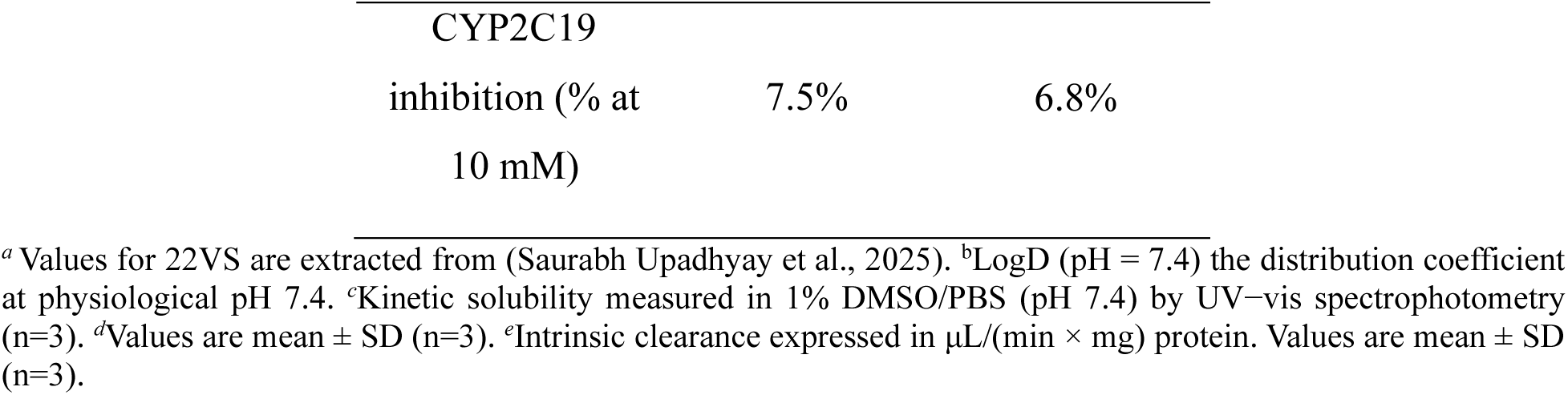
In vitro PK profiles of compounds 22VS and BPU11.

The compound was highly stable in both simulated gastric and intestinal fluids (> 95% remaining after 2 h), suggesting favorable resilience to gastrointestinal metabolism. In microsomal stability assays, BPU11 demonstrated improved metabolic stability relative to 22VS, with mouse t_1/2_ = 78.6 ± 3.9 min and human t_1/2_ = 95.2 ± 4.4 min, corresponding to intrinsic clearance values of 24.8 ± 2.5 µL/min/mg and 18.6 ± 1.7 µL/min/mg, respectively.

BPU11 was also stable in mouse and human plasma (t_1/2_ > 120 min) and in PBS (t_1/2_ > 450 min). Plasma protein binding was high (97.9 ± 0.6%), consistent with a moderate hydrophobic character. The compound exhibited no measurable cytotoxicity against WI-38 or HS-27 fibroblasts (IC50 > 100 µM) and did not inhibit hERG currents up to 80 µM. Moreover, BPU11 showed minimal inhibition of key CYP450 isoforms, with residual activities of > 90% at 10 µM across CYP3A4, 2D6, 2C9, and 2C19 (Table 2).

Collectively, these findings highlight the superior stability, solubility, and metabolic profile of BPU11, supporting its advancement as a preclinical candidate with improved pharmacokinetic properties relative to 22VS.

### 3.7. Evaluation of T Cell Activation in Tumor-PBMC Coculture

We next assessed the immunomodulatory activity of BPU11 in a 3D tumor–PBMC co-culture system that recapitulates CD28-dependent costimulation within a tumor microenvironment. IFN-γ–pretreated A549 tumor spheroids were co-incubated with human PBMCs (effector-to-target ratio = 5:1) in the presence of sub-optimal anti-CD3. Under these conditions, the anti-CD3 only group showed robust T-cell activation, with IFN-γ levels of 145 ± 9 pg/mL (Figure 8A), consistent with CD28-mediated co-stimulation via tumor-expressed CD80/CD86.

**Figure 8.**
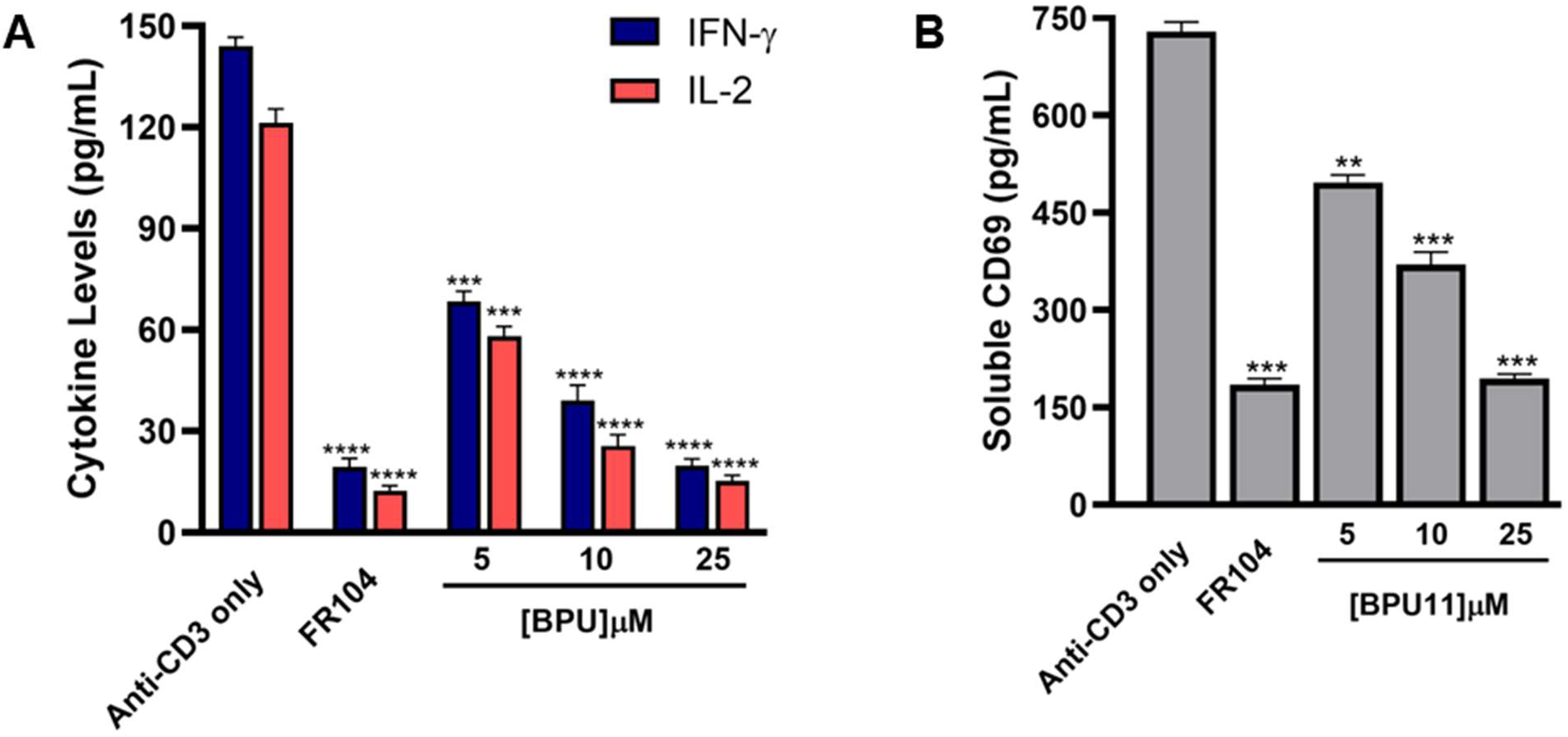
Dose-dependent inhibition of T cell activation markers in tumor–PBMC co-culture. **(A)** Soluble IFN-γ and IL-2 levels were quantified after 48-hour co-culture of A549 tumor spheroids with human PBMCs (E:T ratio 5:1) in the presence of anti-CD3 (0.3 µg/mL) and test compounds: FR104 (10 µg/mL) or BPU11 (5, 10, 25 µM). **(B)** Soluble CD69 levels measured by ELISA after 48-hour co-culture of A549 tumor spheroids with human PBMCs (E:T ratio 5:1) in the presence of anti-CD3 (0.3 µg/mL) and test compounds: FR104 (10 µg/mL) or BPU11 (5, 10, 25 µM). Data represent mean ± SEM of n = 6 independent wells. Statistical comparisons to the “Anti-CD3 only” group were performed using one-way ANOVA followed by Dunnett’s post-hoc test. ** *p* < 0.01, *** *p* < 0.001, and **** *p* < 0.0001 relative to anti-CD3.

As a benchmark, FR104 (10 µg/mL) nearly abolished activation, reducing IFN-γ secretion to 17– 22 pg/mL across replicates (Figure 8A). Treatment with BPU11 led to a clear, concentration-dependent suppression of cytokine release. At 5 µM, IFN-γ levels fell to 71 ± 6 pg/mL; at 10 µM, to 35 ± 5 pg/mL; and at 25 µM, to 18 ± 3 pg/mL (Figure 8A), approaching the magnitude of FR104 inhibition. Similar dose-dependent decreases were observed for IL-2 (Figure 8A) and soluble CD69 (Figure 8B), each declining by >70% at 25 µM relative to anti-CD3 alone.

The potency of BPU11 was thus markedly improved compared to the earlier analog 22VS, achieving near-complete blockade of CD28-driven T-cell activation at one-half the concentration required previously. These data validate BPU11 as an optimized, small-molecule CD28 antagonist capable of functionally mimicking biologic inhibitors within a physiologically relevant co-culture system.

### 3.8. BPU11 Suppresses CD28-Dependent Cytokine Release in a Human PBMC–Mucosal Co-culture Model

To evaluate whether BPU11 maintains CD28-targeted immunosuppressive activity at a physiologically relevant epithelial–immune interface, we employed a human PBMC–mucosal co-culture system. Fully differentiated MucilAir™ airway epithelial tissues were overlaid with primary PBMCs and stimulated with plate-bound anti-CD3 and soluble anti-CD28 for 48 h. Cytokine secretion into the apical compartment was quantified by ELISA (Figure 9).

**Figure 9.**
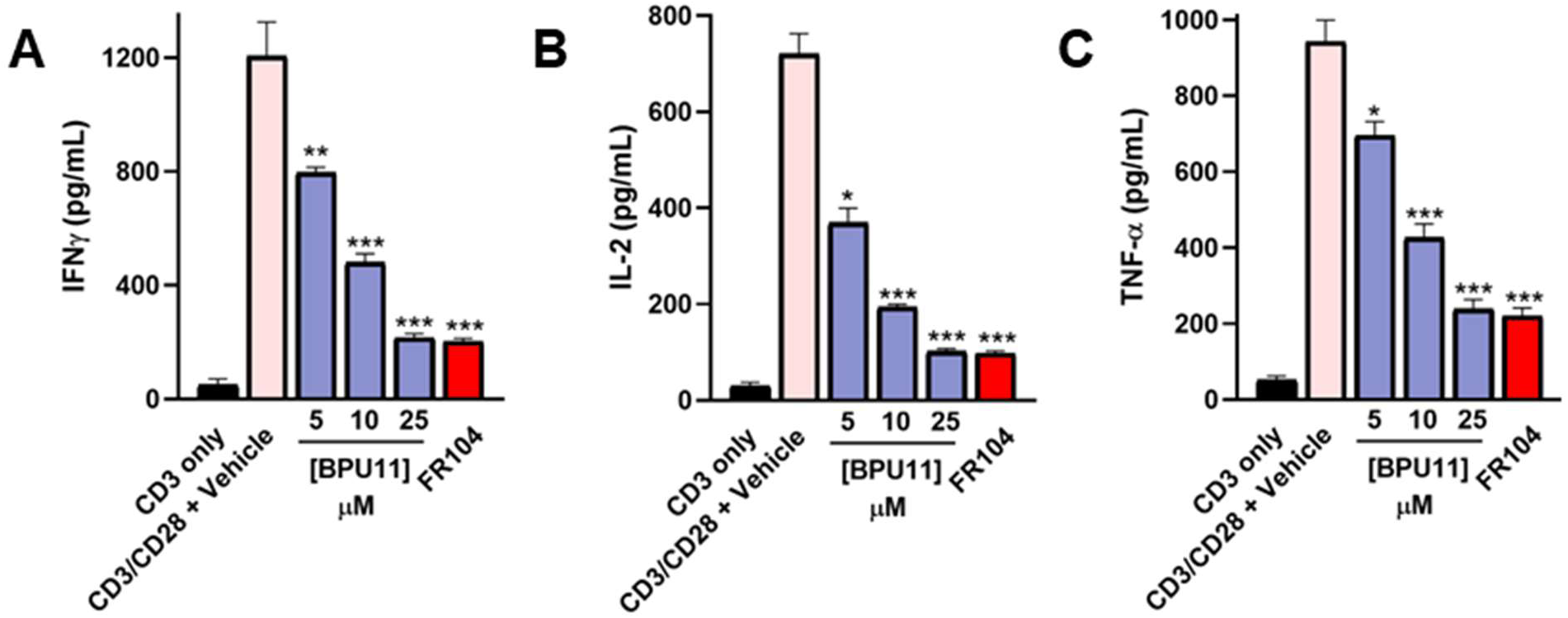
The impact of BPU11 on CD28-dependent cytokine release in a human PBMC– mucosal co-culture model. Quantification of IFN-γ **(A)**, IL-2 **(B)**, and TNF-α **(C)** secretion (pg/mL) in apical supernatants by ELISA. BPU11 suppressed CD28-induced cytokine production in a dose-dependent manner. Statistical comparisons to the “CD3/CD28 + Vehicle” group were performed using one-way ANOVA followed by Dunnett’s post-hoc test. * *p* < 0.05 and *** *p* < 0.001 relative to “CD3/CD28 + Vehicle” group. Error bars represent the standard error mean from n=5 replicates.

As shown in Figure 9, dual stimulation with anti-CD3/CD28 produced robust cytokine release (IFN-γ, IL-2, TNF-α) relative to anti-CD3 alone. The biologic antagonist FR104 (10 µg/mL) potently suppressed this activation, reducing IFN-γ, IL-2, and TNF-α to ∼130 ± 20, 70 ± 12, and 190 ± 25 pg/mL, respectively. BPU11 exhibited clear, concentration-dependent inhibition across all readouts. At 5 µM, IFN-γ and IL-2 were reduced to ∼790 ± 25 pg/mL and 382 ± 18 pg/mL, respectively (Figures 9A and B); TNF-α fell to 692 ± 21 pg/mL (Figure 9C). Increasing the dose to 10 µM further suppressed cytokine levels (IFN-γ ≈ 421 ± 13 pg/mL; IL-2 ≈ 202 ± 11 pg/mL; TNF-α ≈ 211 ± 9 pg/mL). At 25 µM, cytokine output was near baseline, overlapping with the FR104 profile (Figure 9).

These results demonstrate that BPU11 maintains potent and selective suppression of CD28-dependent immune activation in a complex human mucosal tissue model, mirroring its efficacy in the tumor–PBMC assay. The consistency across two independent systems underscores the robustness of BPU11 and supports its advancement as a next-generation, small-molecule CD28 pathway inhibitor.

## 4. Discussion

Resistance to PD-1 blockade and the clinical limitations of CTLA-4 inhibitors are increasingly attributed to persistent CD28 costimulation, which sustains T-cell proliferation even under inhibitory signaling (Wei et al., 2018). Yet, despite its centrality in both antitumor immunity and immune pathology, the CD28–B7 axis has resisted pharmacological modulation outside of biologics (Pulanco et al., 2023). Antagonistic antibodies and CTLA-4-Ig fusions validate the pathway but are constrained by systemic immunosuppression, immunogenicity, and limited tissue penetration (Farhangnia et al., 2023). Here, we provide the first systematic evidence that CD28 can be directly targeted by small molecules through rational scaffold optimization, overturning long-standing assumptions of its “undruggable” nature.

From a structural standpoint, our TRIC and MST screening pipeline identified three 8VS-derived derivatives—BPU11, BPU16, and BPU18—with clear CD28 binding activity. Among these, BPU11 consistently emerged as the most potent, with a dissociation constant of 6.5 μM, nearly five-to ten-fold stronger than its analogues. The SAR trends observed across the library underscore that subtle chemical substitutions at the aryl terminus critically modulate affinity, validating the 1,3,4-thiadiazole–triazole–thioamide core as a privileged scaffold for optimization. Molecular docking and dynamics further revealed that BPU11 engages key residues (His38, Phe93, Lys95, Lys109, Ser110, Asn111) through a combination of hydrogen bonding, hydrophobic contacts, and π–π stacking, while inducing conformational adaptations that stabilize the receptor in energetically favorable states. This induced-fit plasticity not only provides a structural explanation for binding but also demonstrates that CD28 possesses latent druggable pockets, supporting continued investment in rational optimization.

Functionally, these binding events translated into measurable blockade of CD28–B7 interactions. In ELISA assays, BPU11 inhibited CD28–CD80 engagement with an IC_50_ of 18.9 μM, a four-fold improvement over the parent scaffold (71.6 μM), while BPU18 showed intermediate potency (45.2 μM) and BPU16 was weaker (86.7 μM). This rank order closely paralleled binding affinities, highlighting the internal consistency of our biophysical and biochemical data. More importantly, cellular assays established that this activity is not confined to recombinant systems. In CHO-based NanoBit complementation assays, BPU11 inhibited both CD28–CD80 and CD28–CD86 interactions, with maximal inhibition of ∼40–45% and IC_50_ values of 21.9 μM and 38.7 μM, respectively. These results are notable given the natural affinity difference between CD28 and its ligands (Kd ∼4 μM for CD80 vs. ∼20 μM for CD86), and they demonstrate that a single chemotype can perturb both interfaces in living cells.

The translational relevance of this inhibition was confirmed in the CD28 Blockade Bioassay, where BPU11 suppressed downstream T-cell costimulatory signaling with an IC_50_ of 7.9 μM, representing a nearly three-fold gain in potency compared with 8VS (22.8 μM). This pathway-level validation provides the most compelling evidence that BPU11 directly engages CD28 in a biologically relevant context, moving beyond receptor binding to functional suppression of T-cell activation. Importantly, cytotoxicity profiling distinguished BPU11 from related analogues: Jurkat viability remained >85% up to 300 μM, with reductions only at 500 μM, whereas BPU16 and BPU18 were cytotoxic at ≥30 μM. These data establish BPU11 as not only the most potent analogue but also the safest within the tested series, reinforcing its suitability as a chemical starting point.

Beyond its biochemical potency, BPU11 displayed a favorable pharmacokinetic and translational profile. The in-vitro PK data revealed improved solubility, microsomal stability, and plasma persistence compared to 22VS, establishing the scaffold’s suitability for further preclinical development. Functionally, BPU11 translated these properties into robust immunomodulatory activity across two independent co-culture platforms. In the tumor–PBMC spheroid assay, BPU11 produced dose-dependent suppression of IFN-γ, IL-2, and soluble CD69 secretion, achieving near-complete inhibition at 25 µM that mirrored the biologic antagonist FR104. Similarly, in the PBMC–mucosal tissue model, BPU11 attenuated CD28-dependent cytokine release (IFN-γ, IL-2, TNF-α) in a concentration-responsive manner, confirming its efficacy in both tumor-associated and epithelial immune contexts. Together, these findings demonstrate that BPU11 not only engages CD28 at the molecular level but also sustains functional blockade in physiologically relevant environments, underscoring its promise as a chemically tractable alternative to antibody-based CD28 antagonists.

Together, these findings establish BPU11 as the first SAR-optimized small-molecule antagonist of CD28, with consistent activity across binding, biochemical, and cellular assays, and a safety profile superior to related analogues. Beyond advancing a lead scaffold, this work demonstrates that latent druggable pockets exist within the CD28 dimer interface and can be exploited through structure-guided design. The implications extend beyond immunology: our results place CD28 within the growing class of protein–protein interfaces that can be modulated by small molecules, challenging prior dogma and opening avenues for checkpoint modulation beyond antibodies. Moving forward, optimization toward nanomolar potency, receptor selectivity, and in vivo efficacy will be critical, but the present study provides a clear foundation. Comparable exploratory efforts to identify small-molecule modulators of other immune checkpoints such as ICOS, LAG-3, and even CTLA-4 have highlighted the broader feasibility of drugging PPI-driven costimulatory and coinhibitory pathways, reinforcing the generalizability of our approach (Abdel-Rahman et al., 2025; Sobhani et al., 2024; Zhang et al., 2024). To our knowledge, this represents the first reproducible demonstration of small-molecule antagonism of CD28, redefining its druggability and offering a tangible entry point for next-generation immunomodulators in cancer, autoimmunity, and transplantation.

## Acknowledgements

This work was supported by the National Institute of Diabetes and Digestive and Kidney Diseases (NIDDK) under grant number R01DK137299 (PI: Gabr). We would like to thank the Fisher Drug Discovery Resource Center of Rockefeller University (RRID:SCR_020985) for providing access to the Dianthus and Monolith instruments.

## 6. Data Availability

Data is available upon reasonable request.

