## Supporting Information for "Structure-Guided Optimization and Functional Characterization of Small Molecule Antagonists Targeting CD28 Costimulation"

ORCID: 0000-0001-9074-3331

**Running Title:** SAR derived Small-Molecule Modulation of CD28

**Table S1: Structures and vendor codes for all the derivatives evaluated in this study.**

| S. No. | Manuscript Code | Enamine Code | Structure |
| --- | --- | --- | --- |
| 1 | BPU1 | Z17905141 | 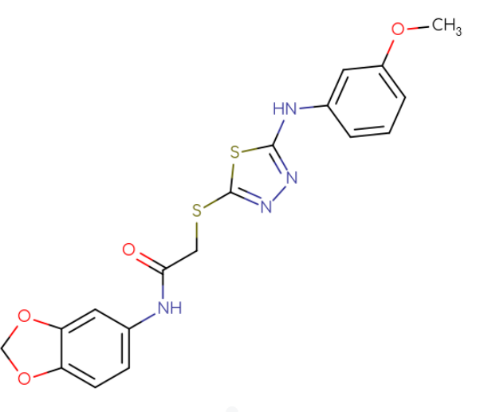 |
| 2 | BPU2 | Z19086246 | 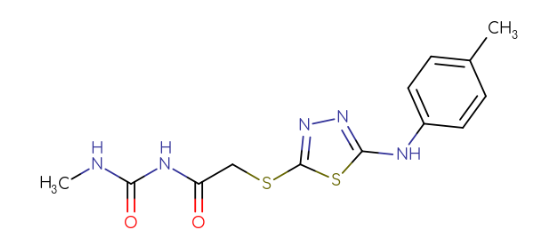 |
| 3 | BPU3 | Z17904608 | 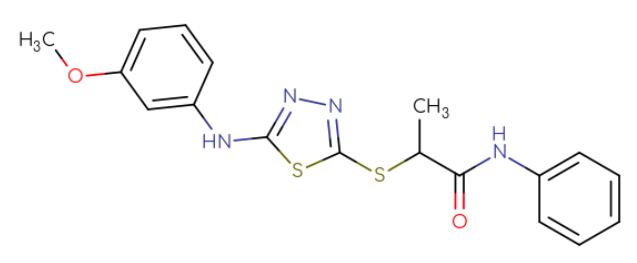 |
| 4 | BPU4 | Z19113595 | 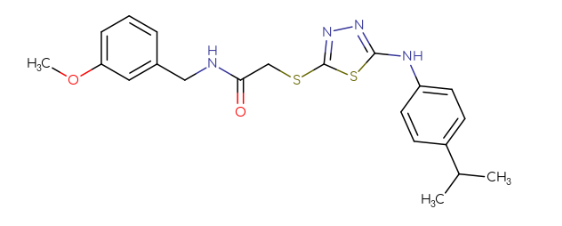 |
| 5 | BPU5 | Z25403700 | 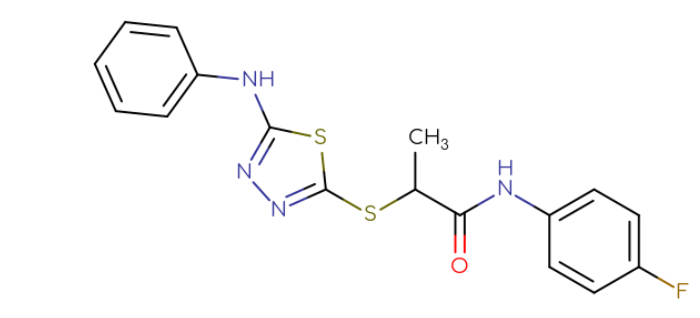 |
| 6 | BPU6 | Z19081223 | 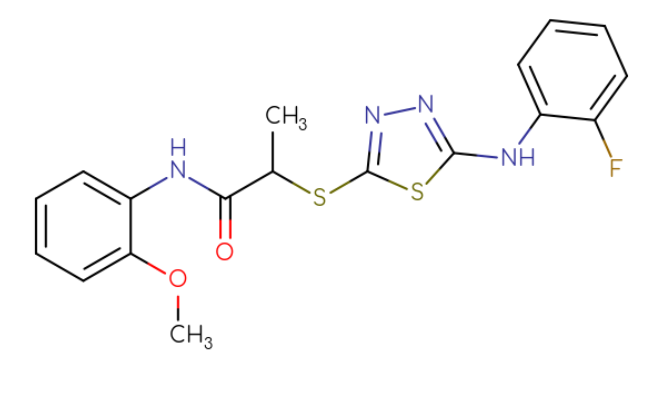 |
| 7 | BPU7 | Z25378486 | 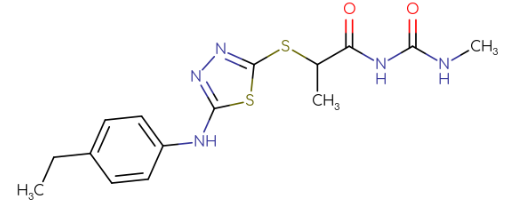 |
| 8 | BPU8 | Z25378522 | 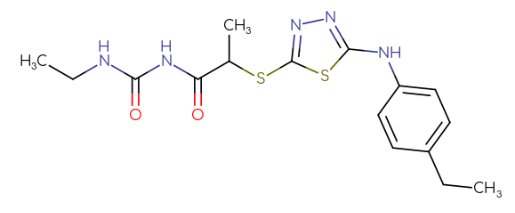 |
| 9 | BPU9 | Z19114077 | 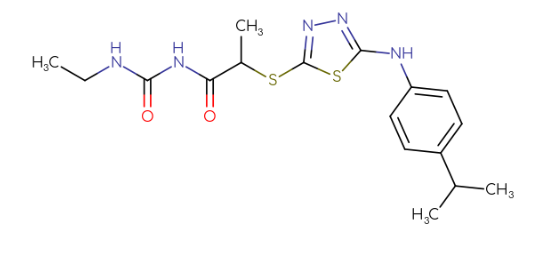 |
| 10 | BPU10 | Z20165405 | 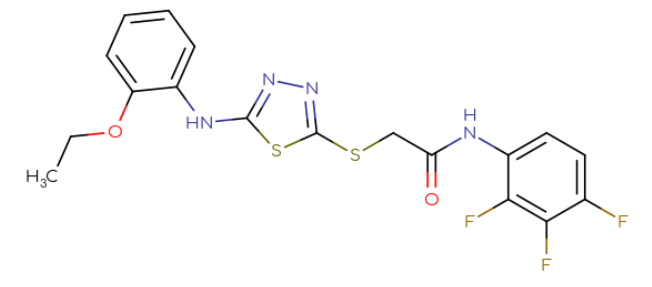 |
| 11 | BPU11 | Z20164101 | 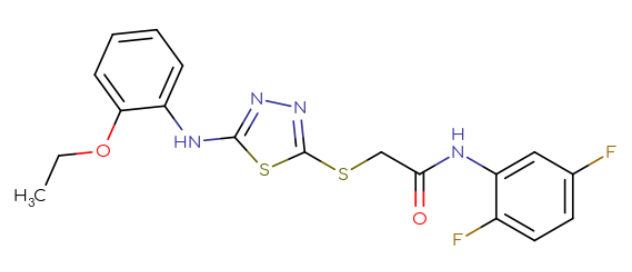 |
| 12 | BPU12 | Z25404068 | 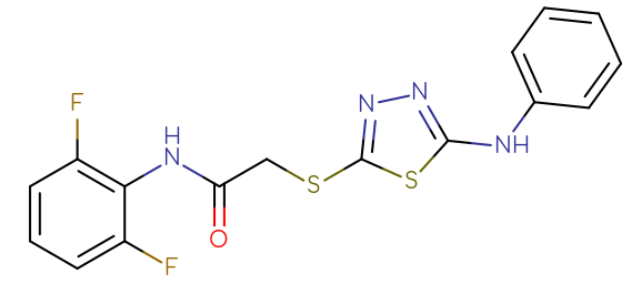 |
| 13 | BPU13 | Z19601415 | 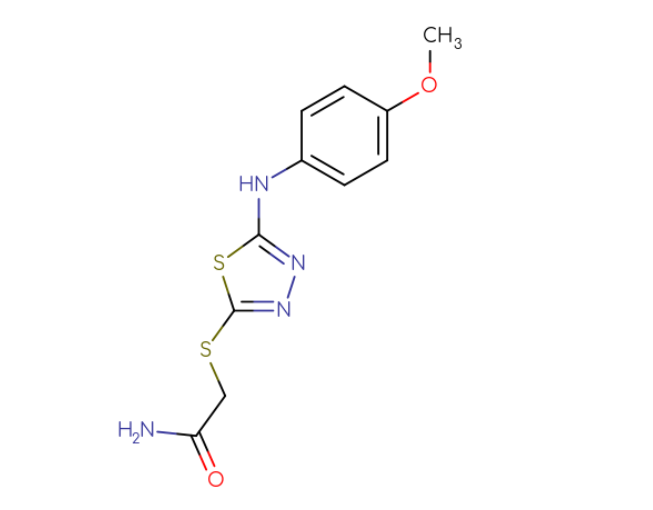 |
| 14 | BPU14 | Z19600740 | 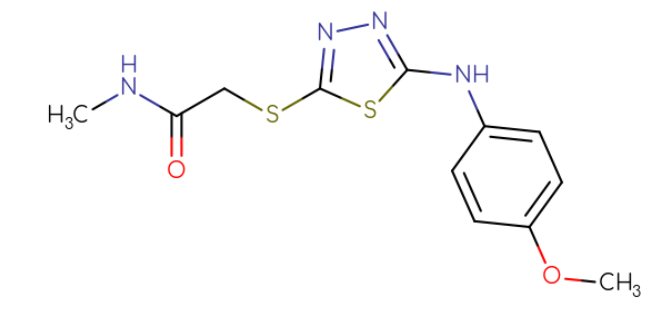 |
| 15 | BPU15 | Z19114449 | 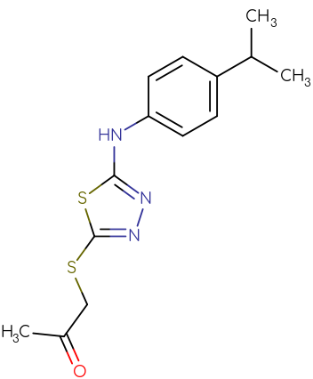 |
| 16 | BPU16 | Z19080488 | 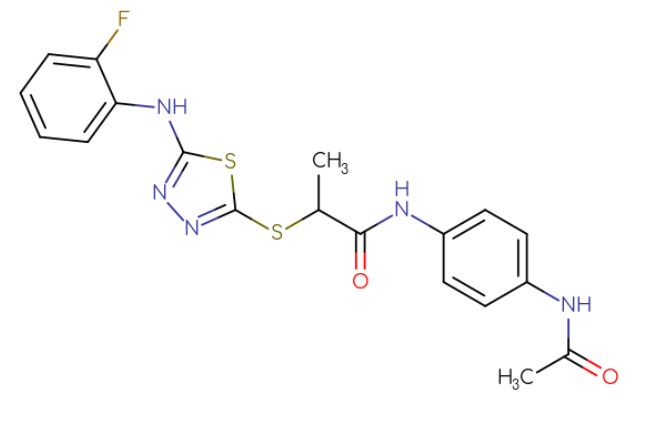 |
| 17 | BPU17 | Z20272487 | 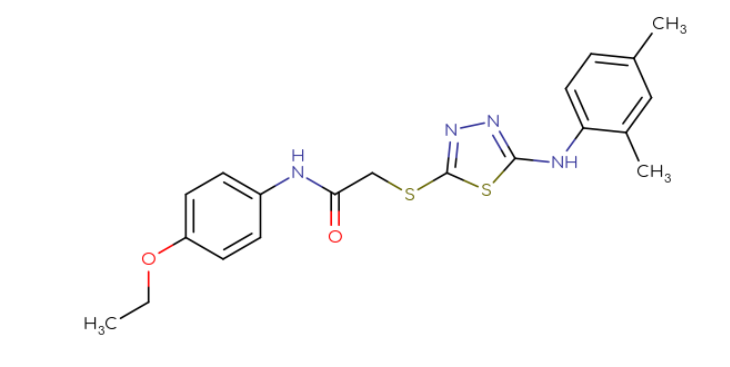 |
| 18 | BPU18 | Z19081686 | 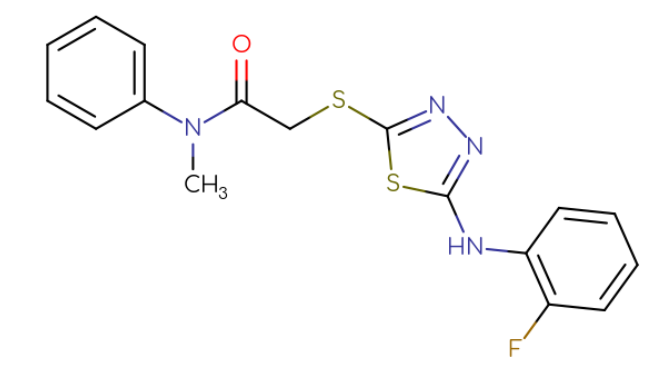 |
| 19 | BPU19 | Z19048228 | 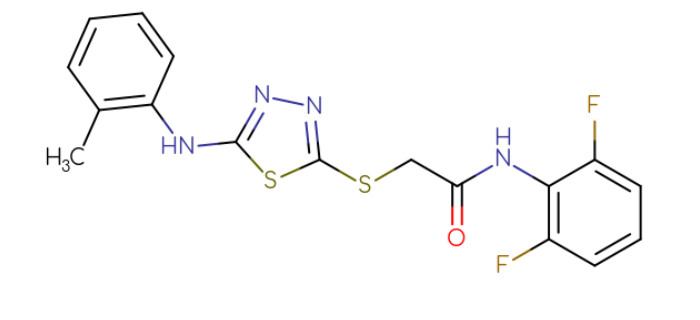 |
| 20 | BPU20 | Z19005948 | 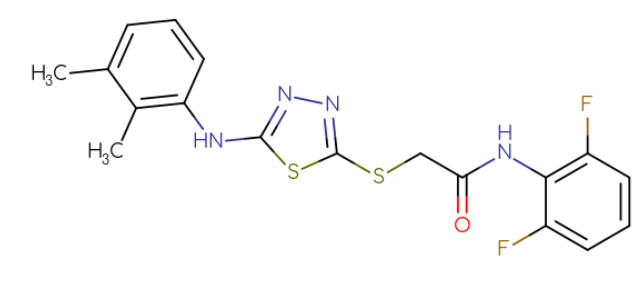 |
| 21 | BPU21 | Z19047008 | 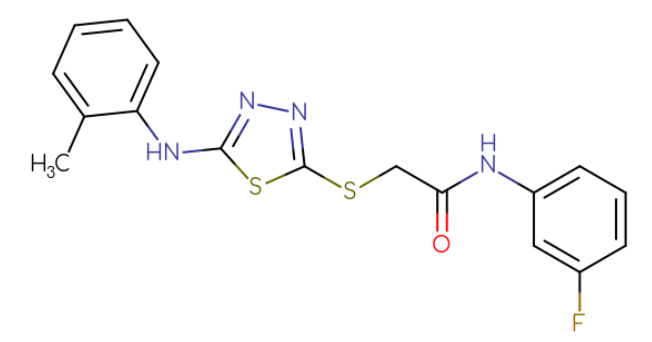 |
| 22 | BPU22 | Z25379089 | 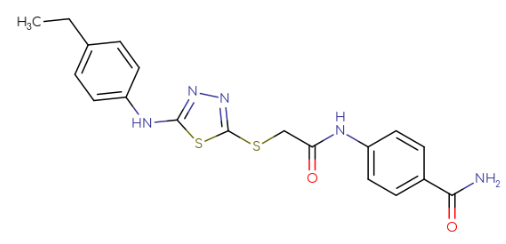 |
| 23 | BPU23 | Z19600745 | 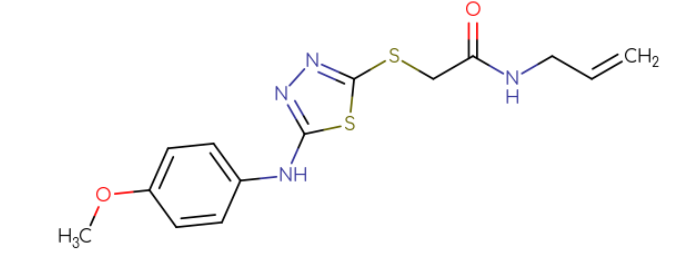 |
| 24 | BPU24 | Z167824220 | 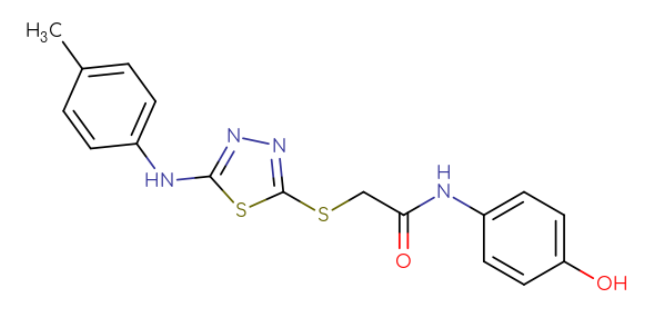 |
| 25 | BPU25 | Z19114134 | 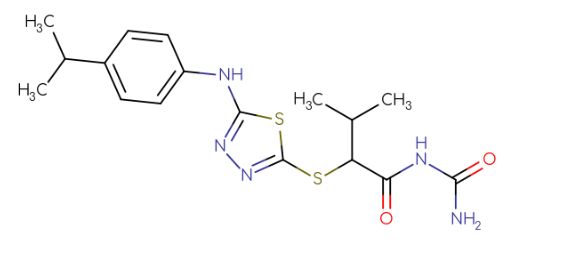 |
| 26 | BPU26 | Z19593565 | 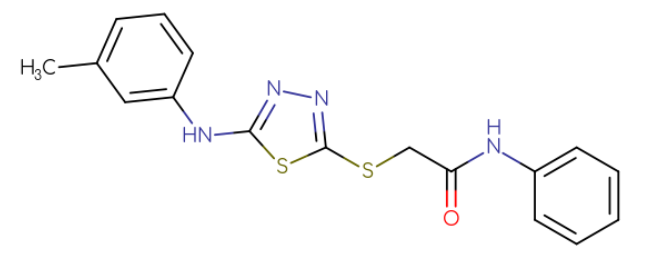 |
| 27 | BPU27 | Z92082494 | 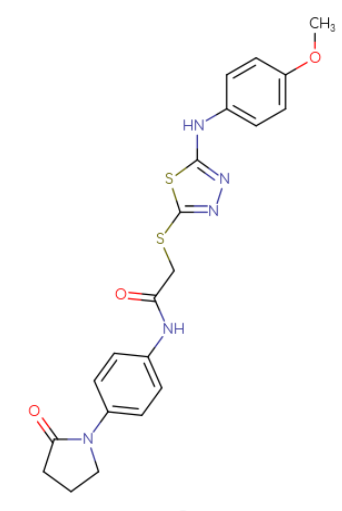 |
| 28 | BPU28 | Z74570151 | 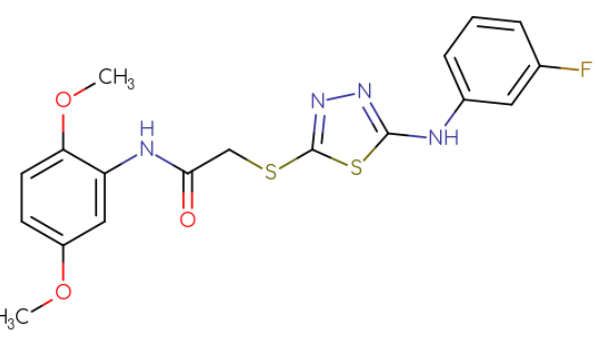 |
| 29 | BPU29 | Z19114407 | 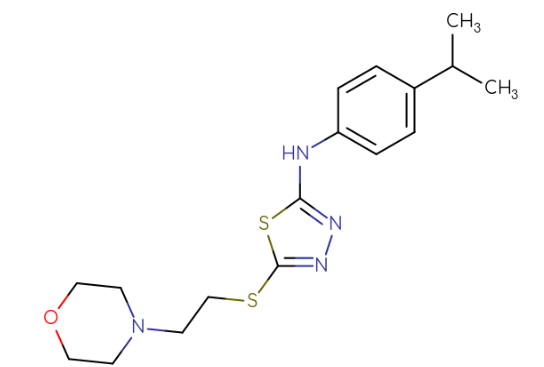 |
| 30 | BPU30 | Z19113845 | 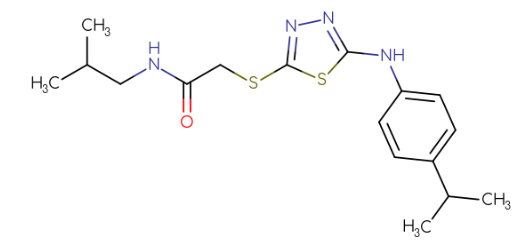 |
| 31 | BPU31 | Z19081718 | 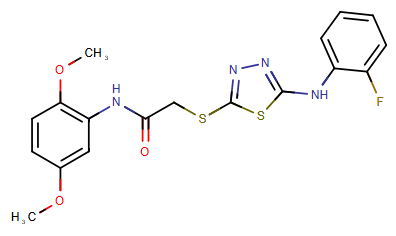 |
| 32 | BPU32 | Z19081750 | 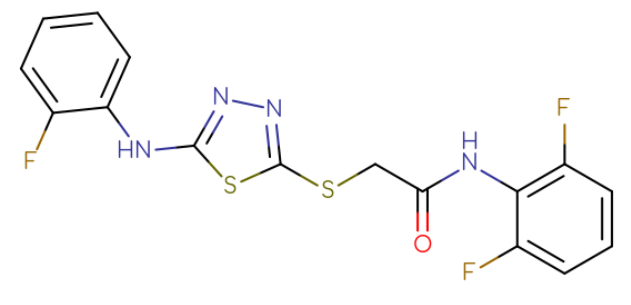 |
| 33 | BPU33 | Z19114797 | 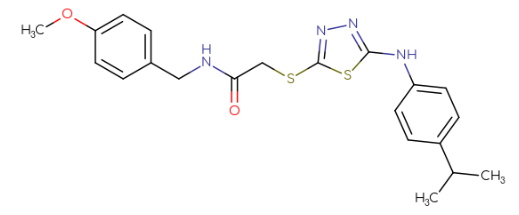 |
| 34 | BPU34 | Z20165446 | 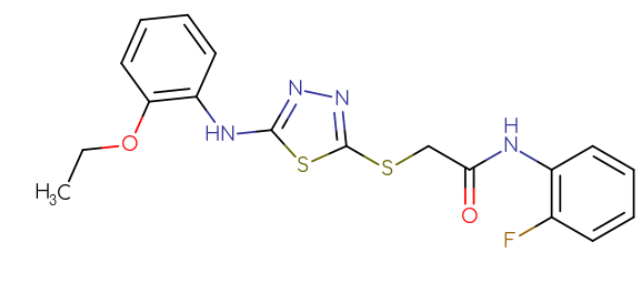 |
| 35 | BPU35 | Z19593490 | 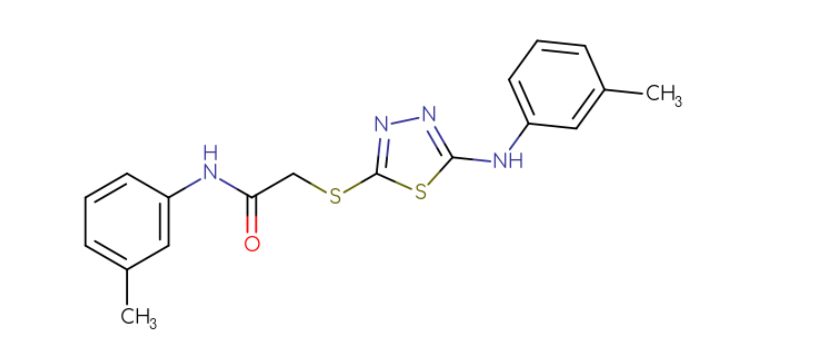 |
| 36 | BPU36 | Z25404048 | 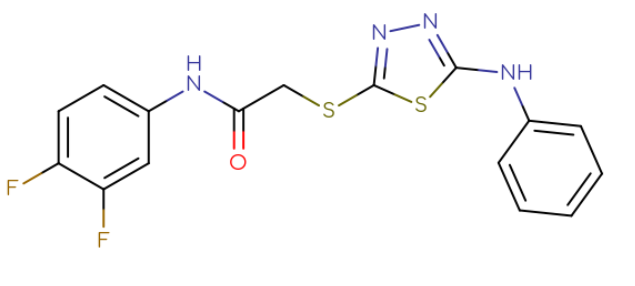 |
| 37 | BPU37 | Z19113855 | 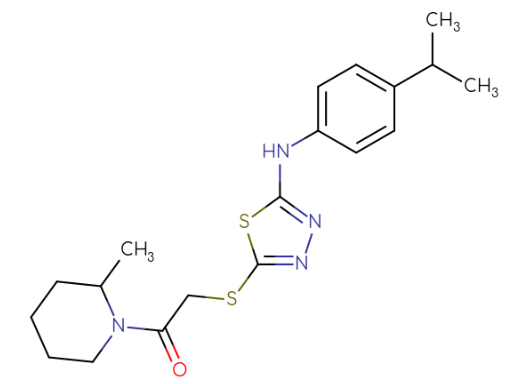 |
| 38 | BPU38 | Z19601621 | 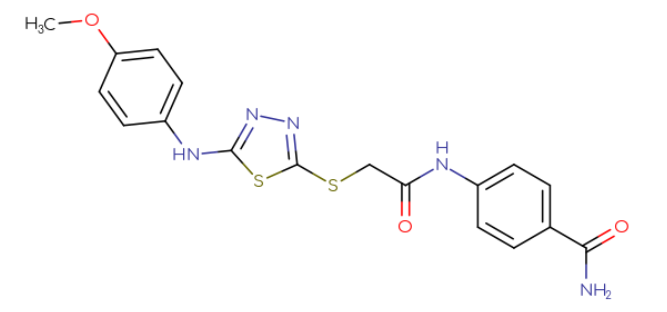 |
| 39 | BPU39 | Z19601651 | 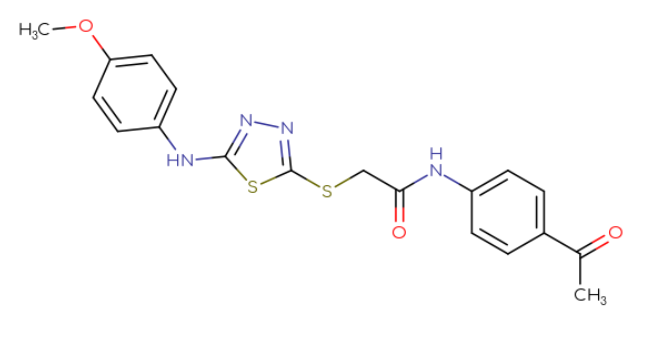 |
| 40 | BPU40 | Z19113766 | 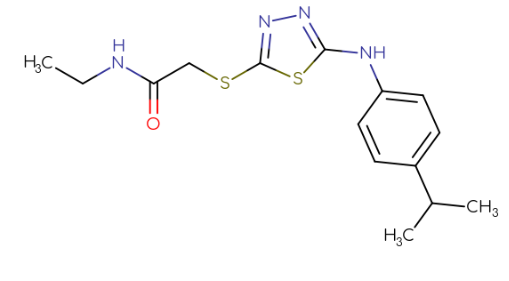 |

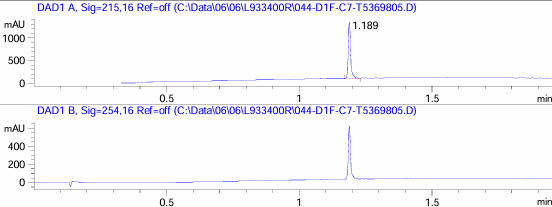

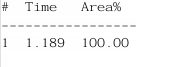

Fig S1: HPLC Spectra for BPU1

Fig S2: HPLC Spectra for BPU2
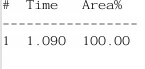

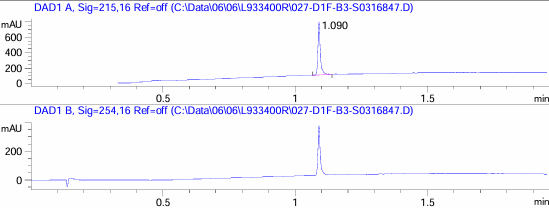

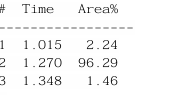

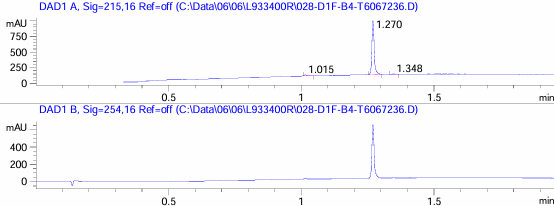

Fig S3: HPLC Spectra for BPU3

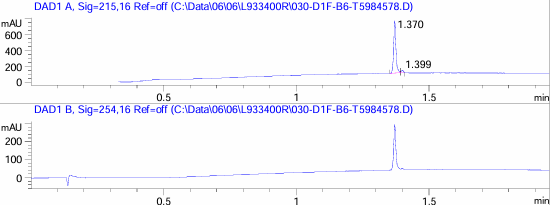

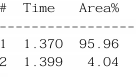

Fig S4: HPLC Spectra for BPU4

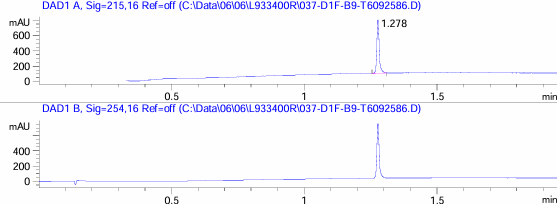

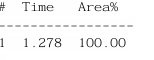

Fig S5: HPLC Spectra for BPU5

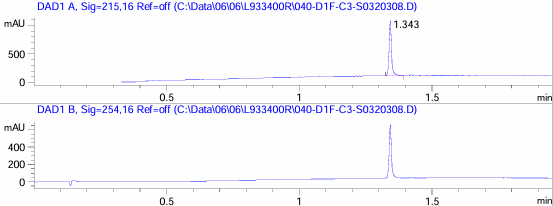

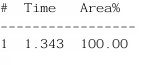

Fig S6: HPLC Spectra for BPU6

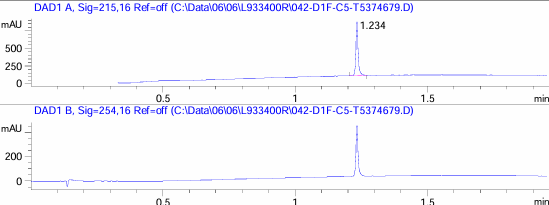

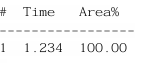

Fig S7: HPLC Spectra for BPU7

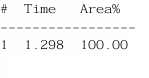

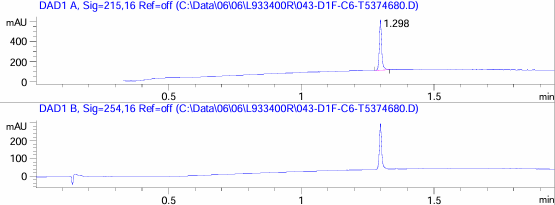

Fig S8: HPLC Spectra for BPU8

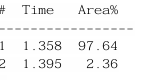

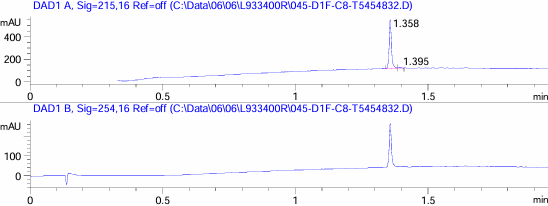

Fig S9: HPLC Spectra for BPU9

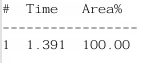

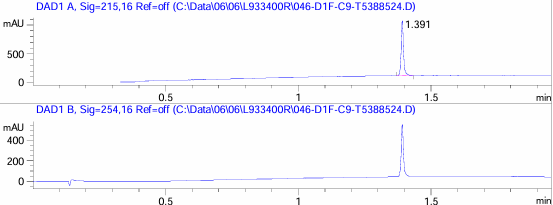

Fig S10: HPLC Spectra for BPU10

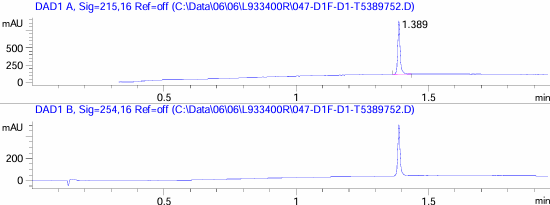

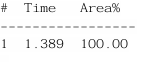

Fig S11: HPLC Spectra for BPU11

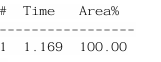

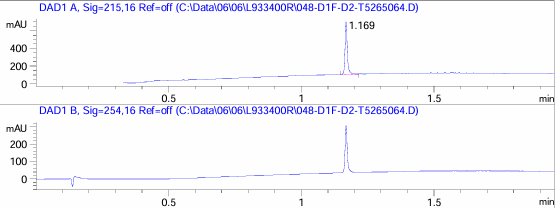

Fig S12: HPLC Spectra for BPU12

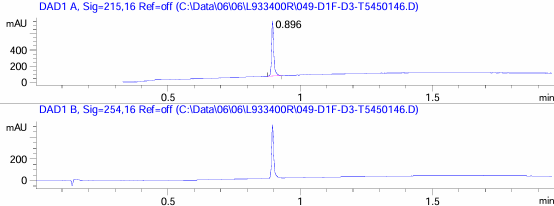

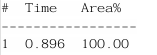

Fig S13: HPLC Spectra for BPU13

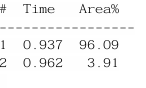

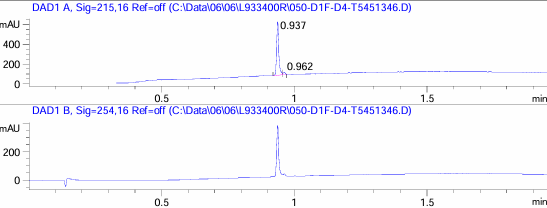

Fig S14: HPLC Spectra for BPU14

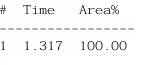

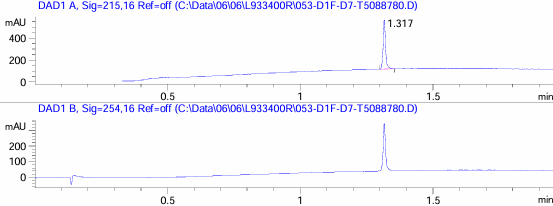

Fig S15: HPLC Spectra for BPU15

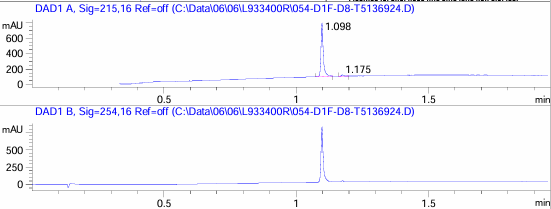

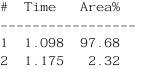

Fig S16: HPLC Spectra for BPU16

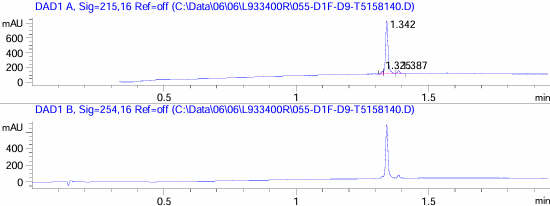

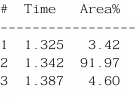

Fig S17: HPLC Spectra for BPU17

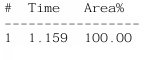

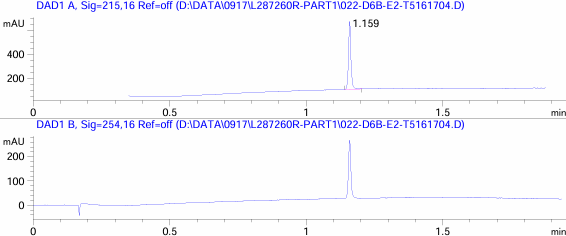

Fig S18: HPLC Spectra for BPU18

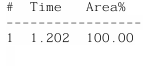

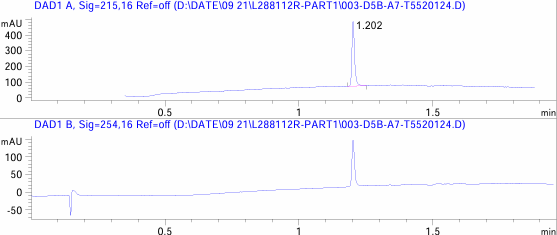

Fig S19: HPLC Spectra for BPU19

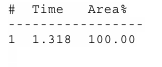

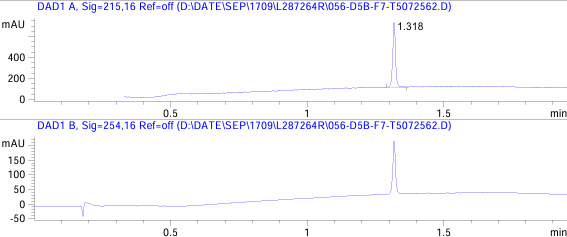

Fig S20: HPLC Spectra for BPU20

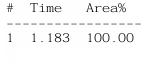

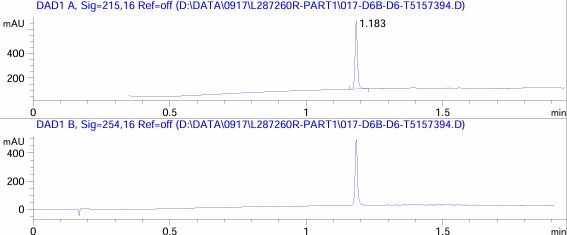

Fig S21: HPLC Spectra for BPU21

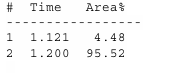

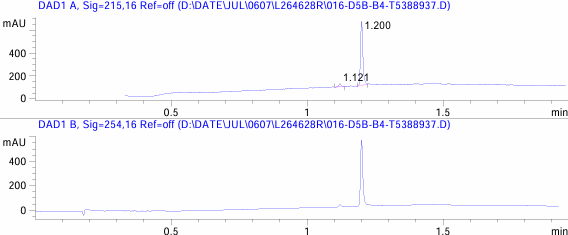

Fig S22: HPLC Spectra for BPU22

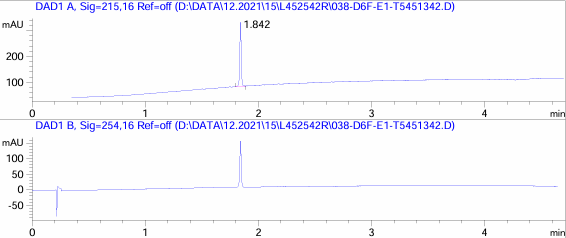

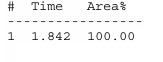

Fig S23: HPLC Spectra for BPU23

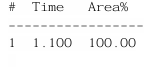

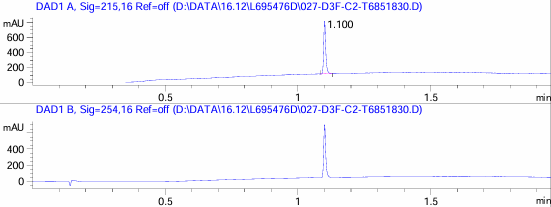

Fig S24: HPLC Spectra for BPU24

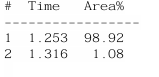

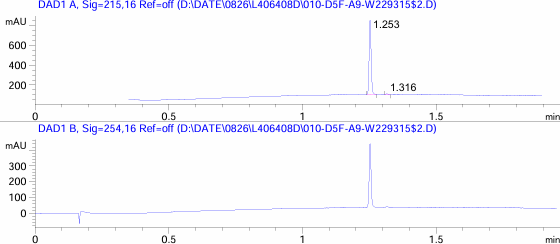

Fig S25: HPLC Spectra for BPU25

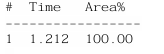

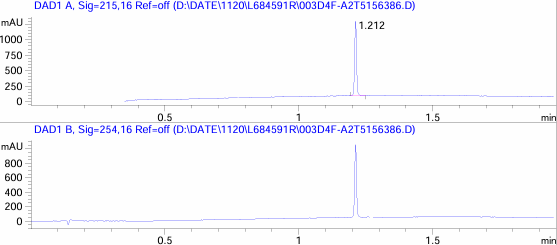

Fig S26: HPLC Spectra for BPU26

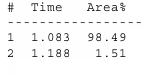

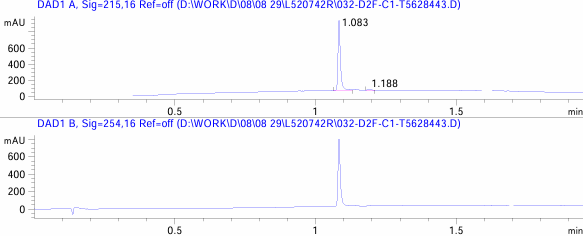

Fig S27: HPLC Spectra for BPU27

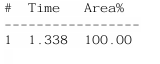

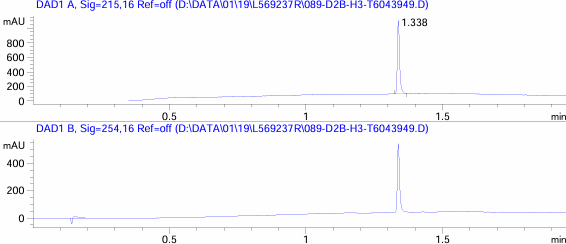

Fig S28: HPLC Spectra for BPU28

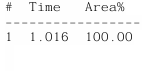

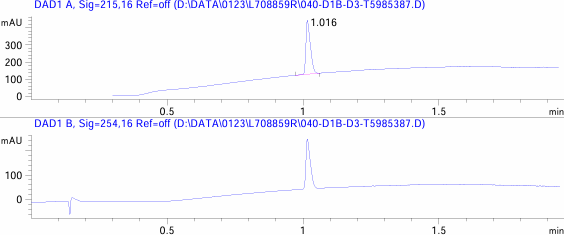

Fig S29: HPLC Spectra for BPU29

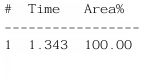

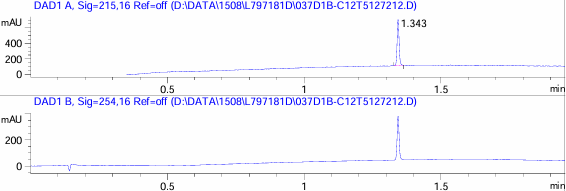
 Fig S30: HPLC Spectra for BPU30

Fig S31: HPLC Spectra for BPU31

Fig S32: HPLC Spectra for BPU32

Fig S33: HPLC Spectra for BPU33

Fig S34: HPLC Spectra for BPU34

Fig S35: HPLC Spectra for BPU35

Fig S36: HPLC Spectra for BPU36

Fig S37: HPLC Spectra for BPU37

Fig S38: HPLC Spectra for BPU38

Fig S39: HPLC Spectra for BPU39

Fig S40: HPLC Spectra for BPU40
